# PacC-dependent adaptation and modulation of host cellular pH controls hemibiotrophic invasive growth and disease development by the rice blast fungus

**DOI:** 10.1101/2020.06.22.164590

**Authors:** Xiao-Lin Chen, Dan He, Changfa Yin, Jun Yang, Jing Sun, Dawei Wang, Minfeng Xue, Zhigang Li, Zhao Peng, Deng Chen, Wensheng Zhao, Jin-Rong Xu, Nicholas J. Talbot, You-Liang Peng

## Abstract

Many of the world’s most serious crop diseases are caused by hemibiotrophic fungi. These pathogens have evolved the ability to colonize living plant cells, suppressing plant immunity responses, before switching to necrotrophic growth, in which host cells die, providing the energy to fuel sporulation and spread of the fungus. How hemibiotrophic pathogens switch between these two lifestyles remains poorly understood. Here, we report that the devastating rice blast fungus, *Magnaporthe oryzae*, manipulates host cellular pH to regulate hemibiotrophy. During infection by *M. oryzae*, host plant cells are alkalinized to pH 7.8 during biotrophic growth, but later acidified to pH 6.5 during necrotrophy. Using a forward genetic screen, we identified alkaline-sensitive mutants of *M. oryzae* that were blocked in biotrophic proliferation and impaired in induction of host cell acidification and necrotrophy. These mutants defined components of the PacC-dependent ambient pH signal transduction pathway in *M. oryzae*. We report that PacC exists as a full-length repressor, PacC^559^, and a truncated transcriptional activator, PacC^222^, which localize to the fungal nucleus during biotrophic growth and to the cytoplasm during necrotrophy. During biotrophy, PacC^222^ directly activates genes associated with nutrient acquisition and fungal virulence, while PacC^559^ represses genes associated with saprophytic mycelial growth and sporulation, which are subsequently de-repressed during necrotrophy. When considered together, our results indicate that temporal regulation of hemibiotrophy by *M. oryzae* requires PacC-dependent sensing and manipulation of host cellular pH.

**Author Summary:** Crop diseases caused by fungi represent some of the most serious threats to global food security. Many fungal pathogens have evolved the ability to invade living plant tissue and suppress host immunity, before switching to a completely different mode of growth, in which they are able to kill host plant cells. This lifestyle– called hemibiotrophy –is exemplified by the blast fungus, *Magnaporthe oryzae*, which causes devastating diseases of rice, wheat and many other grasses. We found that during infection by *M. oryzae*, host cells initially have an alkaline pH, when the fungus is growing in living tissue, but pH rapidly becomes acidic, as host tissue is killed. We identified mutants of the blast fungus that were sensitive to alkaline pH and this enabled us to identify the signal transduction pathway by which the fungus responds to changes in ambient pH. We found that mutants in the pH response pathway were blocked in invasive fungal growth and could not cause acidification of host tissue. Consequently, they are unable to cause blast disease. We characterized the central regulator of this pathway, the PacC transcription factor, which unusually can act as both a repressor and an activator of fungal gene expression. During biotrophic invasive growth, PacC activates many genes previously reported to be required for virulence, including several associated with nutrient acquisition, and at the same time represses genes associated with vegetative growth and sporulation. The PacC signaling pathway is therefore necessary for regulating the switch in fungal lifestyle associated with causing blast disease.

## Introduction

Plant pathogenic fungi can be broadly classified into species that always invade living host tissue, called biotrophs, which evade recognition and suppress host immunity to systemically colonize host plants, and necrotrophic pathogens which overwhelm plant defenses by rapidly killing plant cells, to acquire nutrients from dead or dying tissue [1–3]. Both groups of pathogens exhibit distinct characteristics in terms of the weapons they deploy to infect host plants– such as effector proteins, toxins, and metabolites [4]. Host plants have, in turn, evolved distinct immune signaling pathways to respond to biotrophs and necrotrophs [5]. There is, however, a third group of pathogens, encompassing many of the world’s most serious disease-causing fungi that exhibit both styles of growth. These pathogens are known as hemibiotrophs and initially infect plants like a biotroph, eliciting little response or disease symptoms in their host, but later, at a given point during infection, they switch to killing cells, inducing cellular necrosis and fueling their own sporulation [1]. However, the mechanism by which hemibiotrophic fungal pathogens switch between biotrophy and necrotrophy remains poorly understood [4,6].

Rice blast disease is one of the most devastating diseases threatening rice production worldwide [7–8], and is caused by *Magnaporthe oryzae* (syn. *Pyricularia oryzae*), a hemibiotrophic fungus [6, 9–10]. The fungus initiates infection by forming a specialized infection cell called an appressorium which ruptures the host cuticle allowing invasive hyphae to enter rice cells (IH) [8, 10]. These biotrophic IH are surrounded by a host-derived extra-invasive hyphal membrane (EIHM) [9], and contribute to secretion of apoplastic and cytoplasmic effectors [8, 11]. Apoplastic effectors are often *N*-givcosviated and secreted via the conventional endoplasmic reticulum-Golgi secretion pathway [11–14]. They fulfill diverse roles, including suppression of chitin-triggered immunity [13–14]. By contrast, cytoplasmic effectors are secreted via a plant membrane-derived biotrophic interfacial complex (BIC), using a Golgi-independent process [8, 11–12]. These effectors enable *M. oryzae* to grow in epidermal tissue and move from cell to cell using pit field sites [15]. A fungal nitronate monooxygenase, Nmo2, involved in the nitrooxidative stress response, is also required to avoid triggering plant immunity, thereby facilitating growth and BIC development of *M. oryzae* [16]. Host cells begin to lose viability, once invasive hyphae begin to invade adjacent cells and the switch to necrotrophic growth. This accompanies the appearance of necrotic disease lesions, from which the fungus sporulates [10].

In this study, we set out to explore the mechanism by which *M. oryzae* switches from biotrophic to necrotrophic growth. Specifically, we decided to test the hypothesis that modulation of host cellular pH may be involved in the regulation of this morphogenetic and physiological switch [17]. Alkalinization of plant cells is an important early immune response to attack by pathogens [18–21], and some pathogens have developed mechanisms to alter pH of plant tissues [17, 22]. The necrotrophic pathogens *Athelia rolfsii* and *Sclerotinia sclerotiorum* both, for instance, generate oxalic acid, leading to a sharp drop in the pH of host cells [23–24], while *Fusarium oxysporum* secretes a rapid alkalinization factor to induce host tissue alkalinization [25].

Fungi are generally more acidophilic and have evolved array of mechanisms to adapt to ambient alkaline pH, including the well-known PacC signaling pathway [26]. In *Aspergillus nidulans*, the PacC-dependent pH signaling pathway consists of eight proteins, PacC, PalA, PalB, PalC, PalF, PalH, PalI, and Vps32 [26–27]. The PalH protein is a transmembrane domain-containing protein and functions as an alkaline pH sensor by interacting with an arrestin-like protein, PalF, through its C-terminal domain [28–30]. PalI is also a plasma membrane protein which assists in the localization of PalH [28]. Under alkaline pH conditions, PalF is phosphorylated and ubiquitinated to transduce upstream pH signals [29]. The PalC protein binds to Vps32 by its Bro1-like domain and is a potential linker between the plasma membrane and the endosomal complex [27]. The endosomal protein Vps32 can combine with ESCRT-III (the Endosomal Sorting Complex Required for Transport III) and functions in cargo sorting in the Multi-Vesicular Body (MVB) [31]. The calpain-like protease PalB then assembles Vps32 and PalA as a complex to proteolytically process the transcription factor pacC [32–33].

Under acidic conditions, *A. nidulans* full-length PacC predominantly exists in a protease-inaccessible closed conformation within the cytoplasm. In this conformation, region A (169-301 aa) and region B (334-410 aa) of PacC together interact with the negatively-acting C-terminal domain C (529-678 aa), resulting in a closed conformation [34]. Under alkaline conditions, the interaction is disrupted by upstream Pal proteins and PacC then forms a protease-accessible ‘open conformation’, from which the negatively acting C-terminal domain is removed by two protease cleavage steps. First, the full-length 674-residue pacC72 is cleaved into an intermediate form, pacC53, by removal of the 180 amino acid residues of the C-terminus [35]. In this process, PalA is bound to two YPXL/I motifs beside the 24-residue highly conserved signaling protease box in the C-terminus of PacC, and PalB is likely to be the protease that catalyzes the cleavage [36]. The intermediate pacC53 is then further processed by removal of an additional 245 amino acid residues from the C-terminus to yield a 250-residue form pacC27 [35, 37]. The functional pacC27 contains three Cys2His2 zinc fingers (ZF) and binds to the *cis*-element GCCARG through ZF2 and ZF3 [38–39]. The GCCARG consensus exists in the promoters of genes expressed preferentially under conditions of alkaline ambient pH and also genes repressed at alkaline ambient pH, but expressed preferentially at acidic ambient pH [38]. Although the PacC orthologues in *Saccharomyces cerevisiae* and *Candida albicans* bind to similar *cis*-elements in the promoters of the regulated genes and are dependent on palB for proteolytical processing, their resulting forms and function are distinct from PacC in *A. nidulans*. In *S. cerevisiae*, the PacC counterpart Rim101p is processed by a single step and functions mainly as a transcriptional repressor [40–41]. In *C. albicans*, CaRim101p is an 85kDa protein with a 74-kDa form in alkaline pH and a 65-kDa form at acidic pH, respectively [42]. CaRim101p can also function as both an activator and a repressor. For example, it can directly activate PHR1 (the alkaline-expressed gene) and repress PHR2 (the acidic-expressed gene) [43–44]. However, how fungal PacC proteins function as both transcriptional activators and repressors is still relatively poorly understood.

Here, we show that upon infection by *M. oryzae*, host plant cells are alkalinized to pH 7.8 during the biotrophic growth stage, but then acidified to pH 6.5 at the onset of necrotrophic growth. We report that fungal adaptation to host alkalinization and the induction of host acidification requires the PacC pH signaling pathway. We used a forward genetic screen to identify mutants that were sensitive to alkaline pH, and found that they were all impaired in virulence. We went on to characterise the PacC signaling pathway in the rice blast fungus. We show that in *M. oryzae* the PacC transcription factor simultaneously exists as both a truncated transcriptional activator and a full-length transcriptional repressor, that both localize to the nucleus during biotrophic growth of *M. oryzae*. PacC acts as a key transcriptional regulator which coordinates expression of more than 25% of the protein-encoding genes in *M. oryzae* to facilitate the hemibiotrophic switch, which is necessary for rice blast disease. Plant cellular pH is therefore likely to be a key regulatory signal, that is perceived and modulated by *M. oryzae* to control hemibiotrophic growth.

## Results

### Host plant cells are alkalinized during biotrophic growth but acidified during necrotrophic growth of *M. oryzae*

To investigate the effect of *M. oryzae* infection on host cellular pH, we first established a calibration curve (S1A Fig) to measure cellular pH in barley leaf epidermis using the ratiometric fluorescein-based pH sensitive dye method [45]. Using this method, we observed that pH in the initially infected barley epidermal cells was elevated from pH6.8 to pH7.2 at 12 hours post inoculation (hpi) as appressoria penetrated epidermal cells, and peaked at pH7.8 at 18 hpi when primary infection hyphae had differentiated into bulbous invasive hyphae (IH). However, a decrease to pH7.3 then occurred by 30 hpi as IH rapidly occupied host cells, and then to pH6.5 after 36 hpi when the entire epidermal cell was filled with IH (Fig 1A-1B; S1B Fig). We also measured pH in neighboring plant cells. To our surprise, these uninfected plant cells also became alkalinized by 12 hpi, reaching pH 7.8 by 24 hpi but then became acidic once occupied by *M. oryzae* IH at 48 hpi (Fig 1A-1B). These results show that the pH of host cells is significantly alkalinized during initial infection, but acidified once IH proliferate in host tissue.

**Fig 1.**
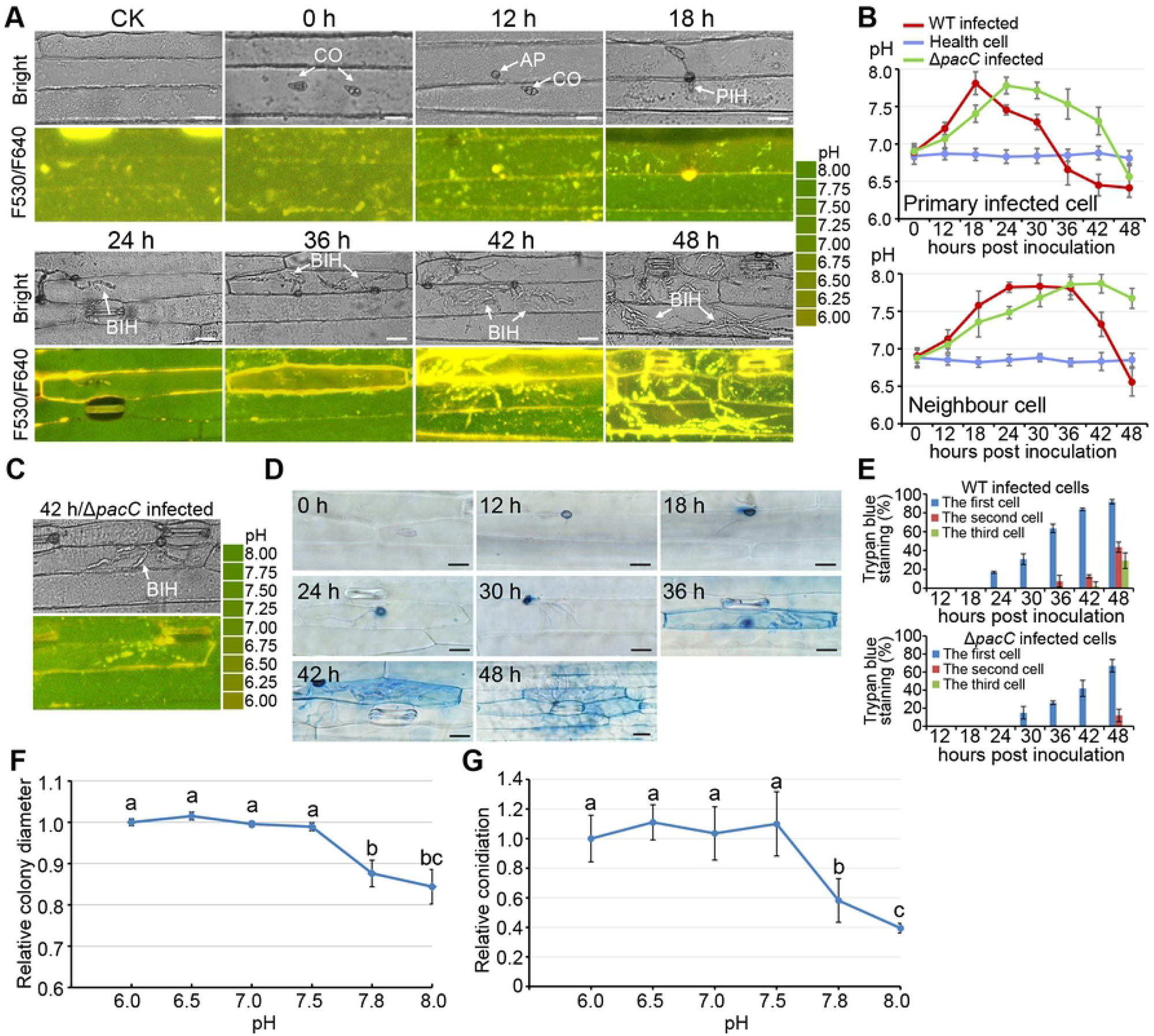
Barley epidermal cells are alkalinized during biotrophic growth but acidified during necrotrophic growth of *M. oryzae*. (A) Bright-field and Ratiometric confocal scanning laser microscopy (CSLM) images of 2’,7’-bis-(2-carboxyethyl)-5-(and-6)-carboxyfluorescein-acetoxymethyl ester (BCECF-AM) stained barley epidermal cells infected with the wild-type *M. oryzae* strain P131 at indicated time points. CO, conidium; AP, appressorium; PIH, primary IH; BIH, branched IH. Bar = 25 μm. (B) Recorded pH levels in barley epidermal cells invaded, (non-invaded) with the wild type and the Δ*pacC* mutant, were calculated by analysing ratiometric CSLM images after BCECF-AM staining at 0-48 hpi. (C) A Bright-field and Ratiometric CSLM image of BCECF-AM stained barley epidermal cells infected with the Δ*pacC* mutant at 42 hpi. (D) Trypan Blue staining of infected plant cells was examined with Nikon90i microscopy at different hpi. Bar = 25 μm. (E) Bar charts showing the percentage of infection sites in which the first, second and third infected cells were stained by Trypan Blue during infection by the wild type strain and Δ*pacC* mutant. (F) Line graph showing inhibition of growth of *M. oryzae* at pH < 5.5 or pH > 7.8. (G) Line graph showing inhibition of conidiation in *M. oryzae* at pH > 7.8. Colony growth and conidiation were measured following growth on complete medium (CM) and oat-broth medium, respectively, and were expressed in relation to the colony diameter or conidiation at pH 6.5.

We then investigated the living viability of infected host cells with Trypan Blue, which selectively stains dead cells [46], and observed that the first infected plant cells at most infection sites only become stained by the Trypan Blue after 36 hpi, (Fig. 1D and E), indicating that initially infected cells are alive before 30 hpi, but lose viability thereafter. Similarly, secondary infected plant cells at most infection sites lost viability after 42 hpi but not before (Fig 1D-1E). Taken together, we reason that host cellular alkalinization occurs during biotrophic growth, probably as an immune response, whereas cellular acidification is associated with the induction of host cell death by *M. oryzae* as the fungus switches to necrotrophic growth.

To understand the physiological consequences of the changes in pH during plant infection, we assayed colony growth and conidiation of *M. oryzae* under different pH. In all three different *M. oryzae* strains assayed, an alkaline of pH 7.8 was inhibitory whereas the acidic pH 6.5 was favorable to both fungal growth and conidiation (Fig 1F; S2 Fig).

### Alkaline-sensitive mutants of *M. oryzae* are impaired in plant infection

To understand the mechanism by which *M. oryzae* adapts to the alkalinized pH of plant cells, we screened a T-DNA insertion mutant library of *M. oryzae* for mutants that were sensitive to pH 7.7 and identified nine mutants (S3A Fig) [47]. The mutants showed reduced colony growth, conidiation and virulence (S3A-3C Fig). Co-segregation analyses indicated that sensitivity to the alkaline pH was caused by the T-DNA insertion (S1 Table). To identify the genes disrupted in these nine mutants, sequences flanking the T-DNA insertion sites were obtained by TAIL-PCR [48]. In mutants CD3179, CD5893, CD9848, and XXY8938, the T-DNA was inserted in promoters or coding regions of *MGG_01615, MGG_06335, MGG_06440*, and *MGG_09311*, respectively. In the remaining mutants, T-DNA was inserted in the promoter or coding region of the same gene *MGG_10150* (S3D Fig). Strikingly, *MGG_01615*, *MGG_06335*, *MGG_06440*, *MGG_09311* and *MGG_10150* are orthologues of *palF*, *palB*, *palH*, *palC* and *pacC* in the PacC-pH signaling pathway in *A. nidulans* [49], and were therefore named *PalF*, *PalB*, *PalH*, *PalC*, and *PacC*, respectively, in this study. Among them, *PacC* encodes the central zinc-finger transcriptional regulator of the pathway.

To confirm mutant phenotypes, targeted gene deletion mutants were generated for *PalF*, *PalB*, *PalH, PalC*, and *PacC* (S4A-4E Fig). All the resulting deletion mutants were sensitive to pH 7.7 and formed darker colonies with reduced growth rates (Fig 2A; S2 Table). In addition, these mutants were reduced in conidiation by 80-90%; compared to the isogenic wild type strain (Fig 2B). In infection assays with barley and rice seedlings, mutants formed tiny lesions mixed with a few larger yellow spots. Under the same conditions, the wild type strain produced numerous larger typical blast disease lesions (Fig 2C). Re-introduction of the corresponding wild-type allele into each null mutant rescued all defects, including sensitivity to alkaline pH. Therefore, disruption of the PacC pH signaling pathway genes makes *M. oryzae* sensitive to alkaline pH and results in a reduction in virulence.

**Fig 2.**
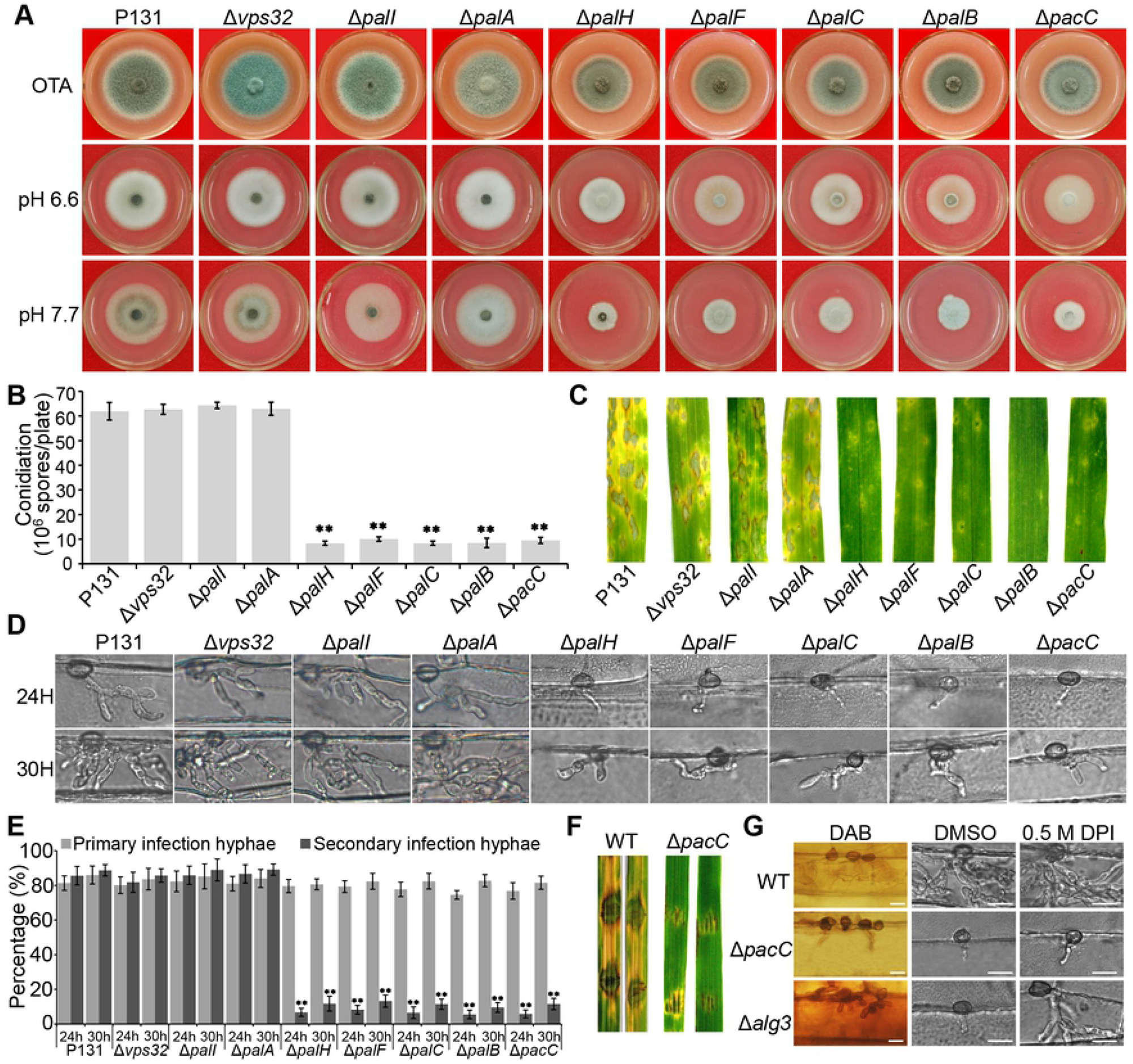
PacC pathway deletion mutants are sensitive to alkaline pH and deficient in biotrophic growth, conidiation and virulence. (A) Colony growth of PacC pathway gene deletion mutants under alkaline pH growth conditions, compared to the wild type (WT) P131. Strains were cultured for 120 h on OTA medium and CM plates at pH 6.6 or 7.7. (B) Bar charts showing conidiation by the same set of strains on OTA plates. Means and standard deviations were calculated from three independent experiments. \**p < 0.01, n* > 100. (C) Blast disease lesions formed on barley leaves by PacC pathway mutants at 5-day post-inoculation (dpi). (D) Micrographs showing development of invasive hyphae (IH) formed by PacC pathway mutants in barley epidermal cells at 24 and 30 h post inoculation (hpi). Bar = 20 μm. (E) Bar chart showing the proportion of appressoria forming primary IH or branched IH at 24 and 30 hpi. (F) Disease lesions formed by the wild type P131 and Δ*pacC* mutant on wounded rice leaves. (G) Micrographs showing that the arrested IH growth phenotype of the Δ*pacC* mutant could not be restored by suppression of host reactive oxygen species (ROS) by diphenyleneiodonium (DPI). A Δ*alg3* mutant was used as the control, which is deleted of the α-1,3-mannosyltransferase gene, induces generation of ROS and is limited in biotrophic growth but can be recovered by addition of DPI (ref. 9). Bar = 20 μm.

In *A. nidulans*, *Pal*I, *Pal*A and *Vps*32 are also involved in the PacC-pH signaling pathway [48]. However, *M. oryzae* mutants disrupted in these three genes were not identified in our screen. We therefore generated Δ*pal*I (MGG_02630), Δ*pal*A (MGG_00833) and Δ*vps32* mutants (S4F-4H Fig). To our surprise, these mutants displayed similar phenotypes to the wild type, including pH sensitivity conidiation and virulence (Fig 2A-2C). Therefore, *PalA, PalI* and *Vps32* are dispensable for regulating the alkaline pH response and virulence, suggesting that these three genes are not required in the PacC signaling pathway of *M. oryzae*, at least during plant infection.

### PacC pathway mutants are impaired in biotrophic growth, induction of host cell acidification and the switch to necrotrophy

To understand why PacC pathway mutants are reduced in virulence, we compared their ability to infect host cells with that of the wild type *M. oryzae* strain P131. During infection of barley leaf epidermis, Δ*palF*, Δ*palB*, Δ*palH*, Δ*palC* and Δ*pacC* mutants showed similar penetration frequencies to the isogenic wild type, but were retarded at the stage of primary infection hyphal growth in more than 70% of infection sites at 24 hpi. Under the same conditions, the P131 developed branched IH at more than 70% of infection sites and by 30 hpi, it had formed branched IH in nearly 90% of infection sites, with invasive hyphae spreading into neighboring plant cells at some infection sites. By contrast, only 10% of appressoria formed branched IH in PacC pathway mutants (Fig 2D-2E). However, Δ*pal*I, Δ*pal*A and Δ*vps*32 mutants were similar to the wild type P131 in development of IH. These data indicate that *M. oryzae* has a PacC pathway that is crucial for biotrophic growth.

To investigate whether subsequent stages of infection are affected by loss of PacC signaling, we inoculated wounded rice leaves to circumvent the need for appressorium-mediated penetration. The wild type P131 generated large lesions, while PacC pathway mutants, including Δ*pacC*, formed significantly smaller lesions without evident necrosis (Fig 2F). We stained Δ*pacC*-infected barley leaves with Trypan blue, and observed that the Δ*pacC* mutant led to host cell death at a much delayed time (Fig 1E), suggesting that the mutant is deficient both in its ability to undertake biotrophic growth and its switch to necrotrophy. In addition, the Δ*pacC* mutant induced less reactive oxygen species (ROS) generation in host cells. However, inhibition of ROS production by diphenyleneiodonium (DPI) treatment failed to allow IH to recover growth (Fig 2G), suggesting that the reduced IH growth of PacC pathway mutants is due to factors other than ROS production.

We also monitored pH changes in barley cells infected by the Δ*pacC* mutant and observed that pH in the initial host cells became alkalinized at 12 hpi, and then peaked at 7.8 at 24 hpi for the initially colonized plant cells and at 36 hpi for the secondary infected cells, respectively (Fig 1B). However, acidification of host cells infected by the Δ*pacC* mutant was much delayed (Fig 1B-1C). These results indicate that the PacC pathway is necessary for inducing host cellular acidification but not for host cellular alkalinization.

### PacC localizes to the fungal nucleus during biotrophic growth and under alkaline ambient pH

To understand how the PacC pathway regulates biotrophic IH growth, we investigated the subcellular localization of the PacC transcription factor during plant infection. We first generated an *eGFP-PacC* fusion construct under control of the native *PacC* promoter and transformed it into a Δ*pacC* mutant (Fig 5B). Subsequent phenotypic assays showed that all the resulting transformants were recovered in colony growth, conidiation and virulence (Fig 5G-5H), indicating that the GFP-fused PacC is functional. Interestingly, GFP signals were predominantly localized to the nucleus in primary and branched IH from 18 to 30 hpi, but were then mainly distributed in the cytoplasm from 36 hpi onwards (Fig 3A). Furthermore, GFP signals reappeared in the nuclei of IH that had penetrated neighboring cells between 36 and 42 hpi, and then disappeared again from the nucleus at 48 hpi (Fig 3A). By contrast, in pre-penetration stage structures, conidia and appressoria, GFP signals were evenly distributed in the cytoplasm (Fig 3A). These data reveal that PacC localizes to the nucleus specifically during biotrophic invasive growth.

**Fig 3.**
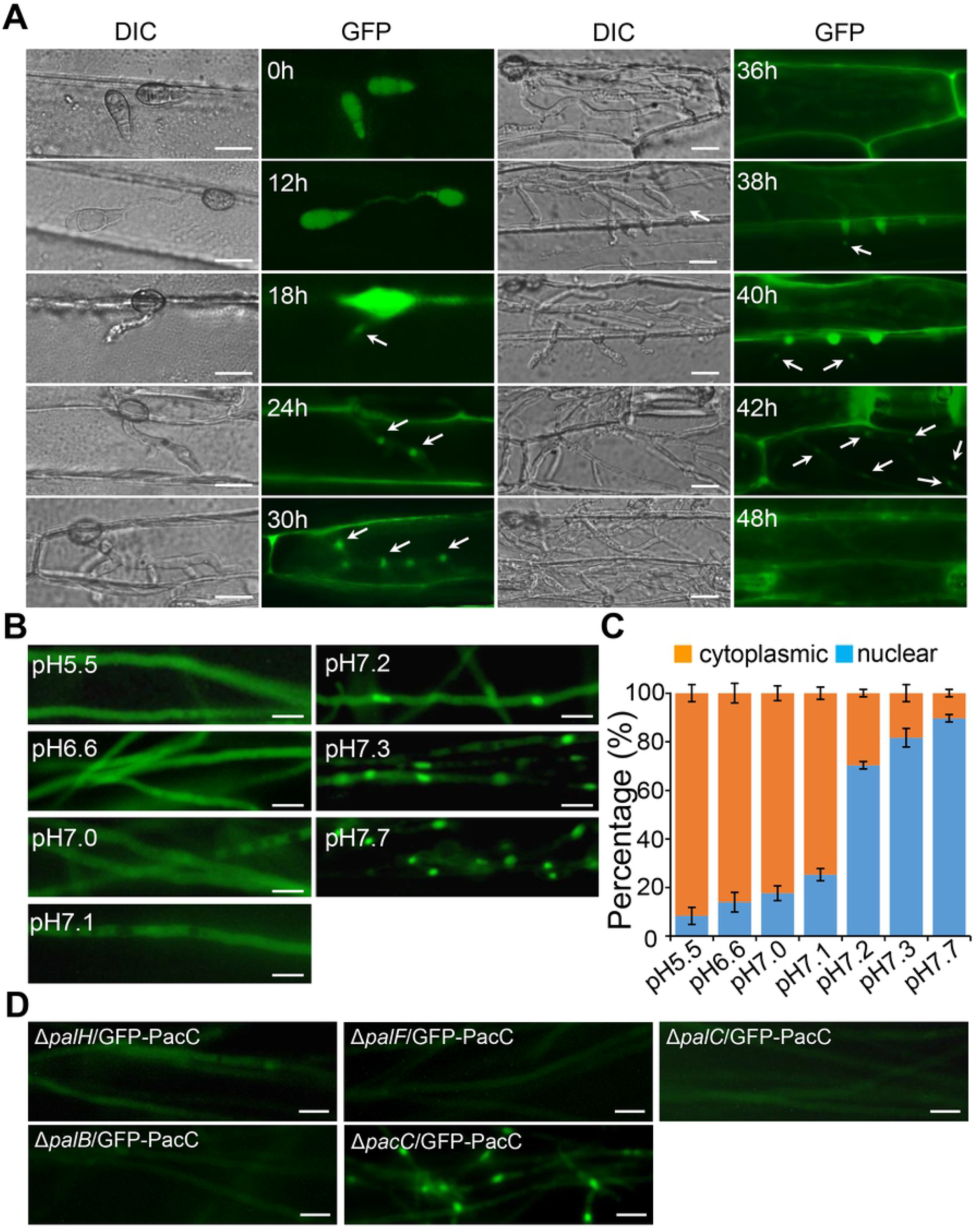
PacC Shows Distinct Subcellular Localization during Biotrophic and Necrotrophic Growth. (A) Bright-field and epifluorescence microscopy micrographs showing that PacC-GFP localizes to the nucleus during biotrophic growth, but to the cytoplasm during necrotrophic growth and asexual development. (B) Micrographs showing that alkaline pH > 7.2 induces nuclear localization of PacC-GFP in the fungal mycelium. (C) Percentage of localization of PacC-GFP in fungal mycelium cultured in different pH conditions. (D) Nuclear localization of PacC-GFP requires the upstream *Pal* genes. PacC-GFP was introduced into Δ*palH, ΔpalF, ΔpalC, ΔpalB* mutants and transformants grown at alkaline pH were visualized by epifluorescence microscopy. Photos were recorded with a Nikon90i epifluorescence microscopy. Bar = 25 μm.

**Fig 4.**
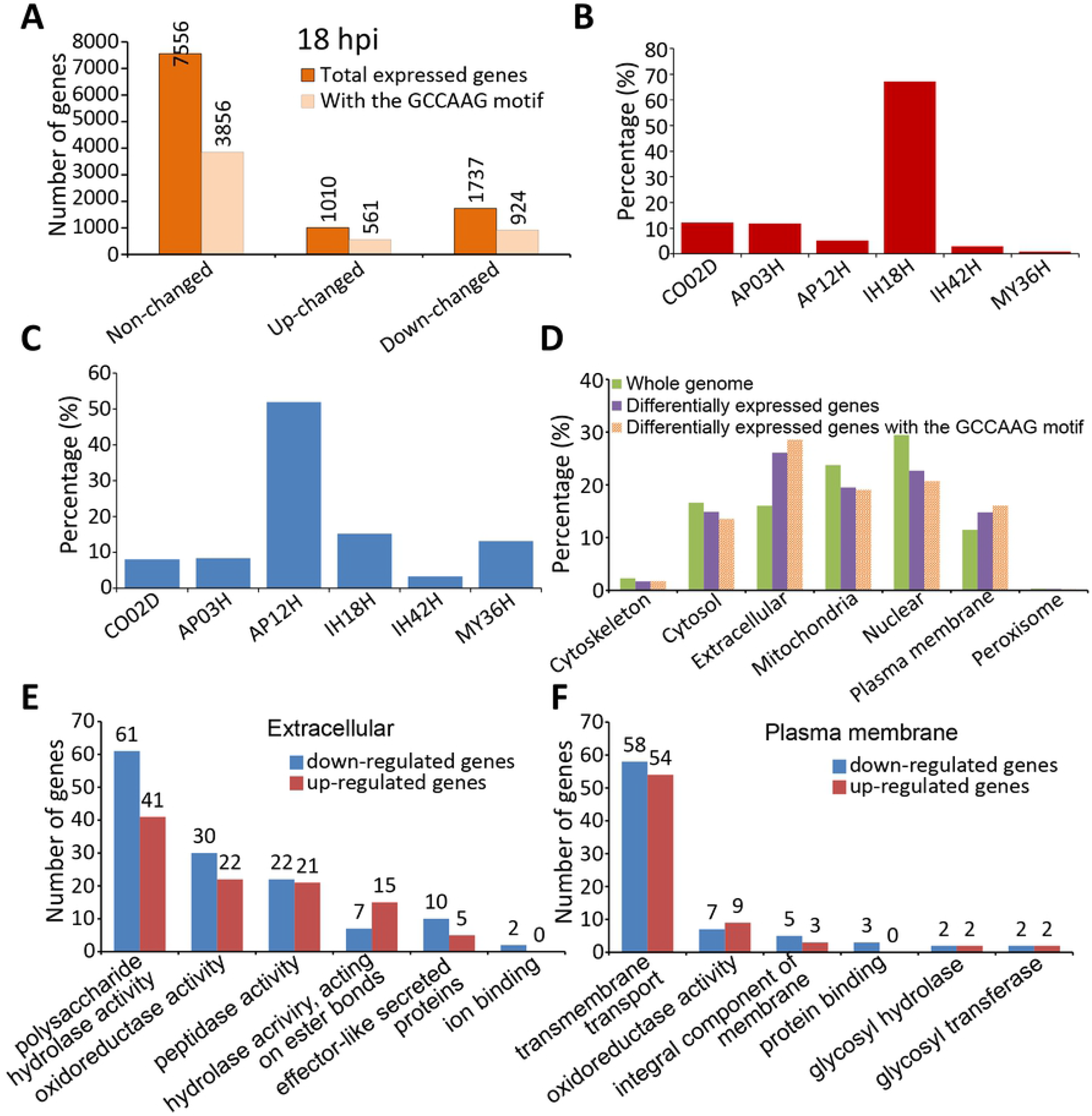
Global Transcriptional Profiling Reveals Major Families of PacC-Regulated Genes. (A) Bar charts showing the number of genes that are altered in expression in a Δ*pacC* mutant compared to the wild type P131, in invasive hyphae (IH) at 18 hpi. Genes containing one or multiple PacC-binding consensus GCCAAG in their promoters are indicated and regarded as PacC-directly regulated genes. (B) Bar charts showing the highest expression stages of the PacC-directly activated genes. (C) Bar charts showing the highest expression stages of the PacC-directly repressed genes. (D) Bar charts showing the predicted subcellular localization of proteins encoded by PacC-directly regulated genes in IH at18hpi. The bars marked ‘Whole genome’ indicates the proportion of genes in the whole *M. oryzae* genome showing each predicted sub-cellular localization pattern. (E) Bar charts showing the predicted functional annotations of putative extracellular proteins encoded by PacC directly regulated genes in IH. (F) Bar charts showing functional annotations of predicted plasma membrane proteins encoded by PacC directly-regulated genes in IH. Two independent duplicates of RNA-seq data were obtained, and genes reproducibly altered in their expression with Log2 > 0.5 or < −0.5, *p* value < 0.005 in the mutant were recorded as PacC-up or down regulated genes.

**Fig 5.**
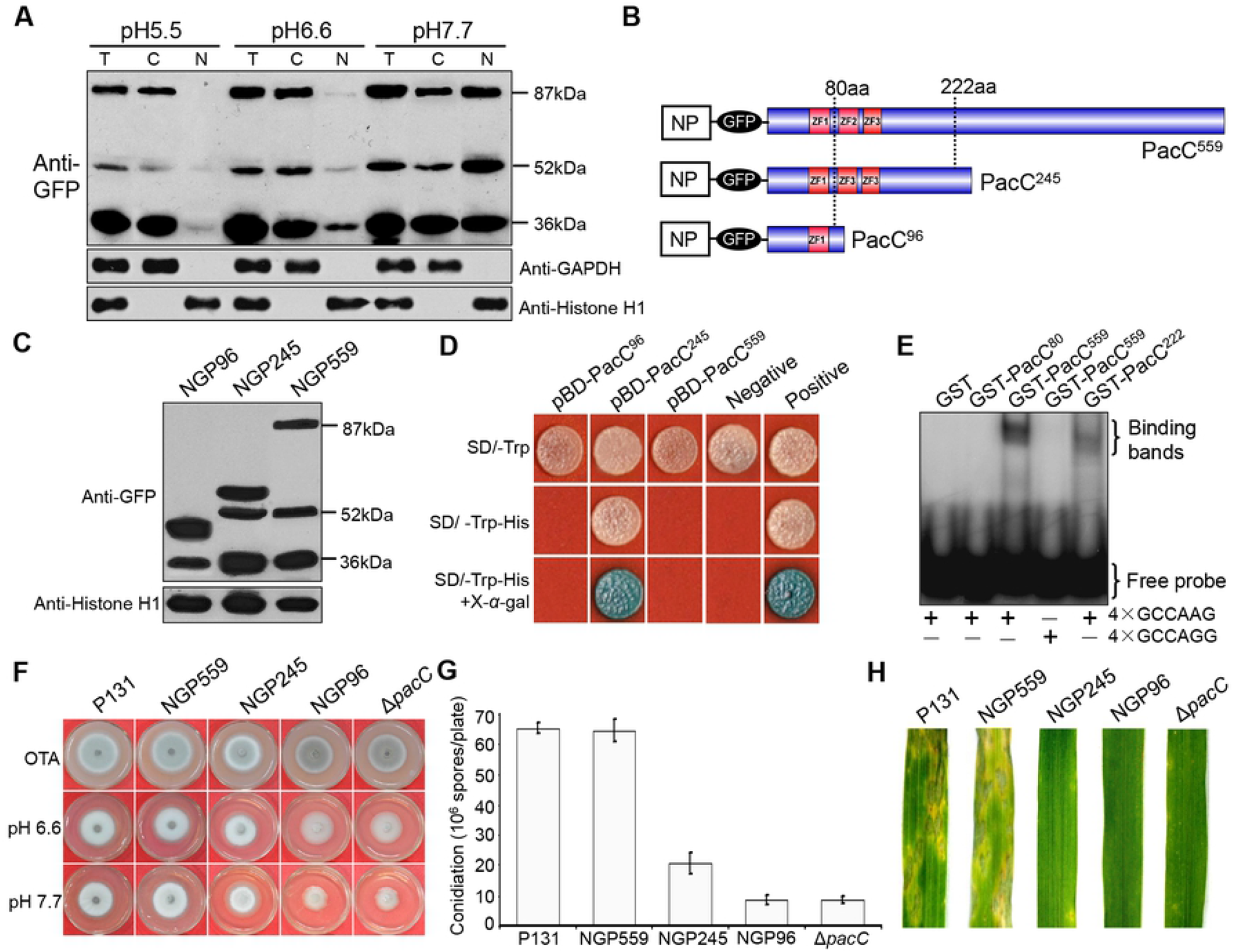
PacC Exists as Both a Full-length Transcriptional Repressor and a Truncated Transcriptional Activator. (A) Western blot showing distinct PacC isoforms detected with anti-GFP antibody in total (T), cytoplasmic (C) and nuclear (N) proteins extracted from mycelium of NGP559 cultured at the pH indicated. Detection using an anti-GAPDH and an anti-histone H1 antibodies was used as the loading control for cytoplasmic and nuclear proteins, respectively. (B) Schematic representation of GFP-PacC constructs transformed into the Δ*pacC* mutant. Dotted lines indicate two putative cleavage sites. NP, native *PacC* promoter; ZF, zinc finger domains. (C) Western blot showing PacC isoforms detected with an anti-GFP antibody in nuclear proteins of transformants NGP96, NGP245 and NGP559 cultured at pH 7.7. (D) Yeast transcription activation assay showing pBD-PacC^559^, pBD-PacC^245^ and pBD-PacC^96^ grown on SD-Trp-His plates and β-galactosidase activity on SD-Trp-His plus X-gal plates. pGBT9 and pGBKT7 were the positive and negative controls, respectively. (E) Electrophoretic mobility shift assay of GST-PacC559, GST-PacC222 and GST-PacC80 showing potential binding to a ^32^P-labelled 4×GCCAAG consensus sequence. (F), (G) and (H) show colony growth by the wild type P131, the Δ*pacC* mutant, and transformants NGP96, NGP245 and NGP559 on OTA and CM at the indicated pH, conidiation of the same strains on OTA and blast disease assays.

Because host cells are alkalinized and acidified during biotrophic and necrotrophic growth, respectively, we suspected that host cellular pH was the inductive signal for PacC nuclear localization. To test this idea, we investigated the subcellular localization of PacC in mycelium cultured under different pH. When the Δ*pacC*/*GFP-PacC* strain was cultured in liquid CM with pH < 7.2, GFP-PacC fluorescence was mainly observed in the cytoplasm (Fig 3B-3C). By contrast, when hyphae were cultured at pH ⩾ 7.2, a proportion of the GFP signal localized to the nucleus; in particular, the majority of the GFP signals localized to the nucleus at pH 7.7, as observed in the branched IH. These observations indicate that alkaline pH is an important signal to induce nuclear localization of PacC, as previously reported in *A. nidulans* [37, 39].

We also introduced the *eGFP-PacC* construct independently into the Δ*palH*, Δ*palF*, Δ*palC* and Δ*palB* mutants, and examined GFP subcellular localization. GFP-PacC was evenly distributed in the cytoplasm, but not in the nucleus of Δ*palH*, Δ*palF*, Δ*palC* and Δ*palB* transformants (Fig 3D). Therefore, the alkaline pH-induced nuclear localization of PacC requires *PalB, PalC, PalF* and *PalH*.

### PacC is a central regulator of gene expression for biotrophic growth of *M. oryzae*

To identify PacC-regulated genes, we then carried out RNA-seq analysis of barley seedlings infected with either the wild type or Δ*pacC* mutant at 18 hpi. The two strains expressed a total of 10303 genes (⩾2 FPKM), of which, 2747 genes were differentially expressed (S1 Dataset), including 1485 genes that have one or multiple GCCAAG *cis*-elements [38] for PacC binding in their promoters (Fig 4A; S1 Dataset) (arbitrarily defined as the 1.5 kb fragment upstream of the translation codon). Among these, 1485 PacC-directly regulated genes, 924 and 561 were repressed or activated in the mutant, respectively (S1 Dataset), suggesting that PacC acts as both a transcriptional activator and a repressor. Interestingly, most of the down-regulated genes showed the highest expression in biotrophic IH (Fig 4B; S2 dataset) whereas most of the 561 up-regulated genes showed the highest expression in penetrating appressoria, as well as during conidiation, germinated conidia or/and mycelium (Fig 4C; S3 dataset), suggesting that during biotrophic growth, PacC enhances biotrophy-related genes and represses genes involved in appressorial penetration, conidiation, and necrotrophic growth.

To define the gene repertoire directly regulated by PacC during biotrophic growth, we analyzed predicted subcellular locations (http://www.genscript.com/wolf-psort.html), protein domains (Pfam, http://pfam.xfam.org/), and gene ontology functions (GO, http://www.geneontology.org/) of their encoded proteins. This analysis revealed that extracellular and membrane proteins were highly enriched (Fig 4D; S4 Dataset). Among predicted extracellular proteins, 255 proteins have Pfam and/or GO annotations, including 172 proteins that are putative polysaccharide hydrolases, peptidases/proteases or lipases (Fig 4E; S4 Dataset). Notably, approximately half of the polysaccharide hydrolases observed are likely to be involved in plant cell wall degradation (S5 Dataset) [50], and the rest may be involved in remodeling the fungal cell wall. In addition, 180 of the predicted membrane proteins have Pfam and/or GO annotations, including 114 putative membrane-associated transporters, which may function together with extracellular hydrolases to acquire carbon and nitrogen sources for biotrophic growth of IH (Fig 4F; S4 Dataset). Furthermore, among genes directly regulated by PacC, over 80 have been functionally characterized and have roles in suppressing plant immunity, cell wall remodeling, nutrient utilization and metabolic processes, including *ALG1, BAS4, BUF1, ECH1, GEL1, GLN1, HEX1, MET13, NMO2, NMR3, PMC2, MoABC7, SDH1, MoAbfB, MoARG7, MoALR2, MoCDA1, MoCDA3, MoCDA4, MoIMD4, MoLDB1, and MoMyo2* (S4 Dataset). When considered together, these data suggest that PacC coordinates gene expression to facilitate biotrophic growth of the fungus and temporally regulates the onset of necrotrophy.

### The PacC transcription factor exists both as a transcriptional activator and repressor

To reveal how PacC simultaneously acts as both a transcriptional activator and a repressor, we performed immunoblot analyses of the eGFP-PacC protein in mycelium of a Δ*pacC*/*GFP-PacC* strain. A full-length 87 kDa fusion protein was detected with an anti-GFP antibody, together with two truncated forms of 52 kDa and 36 kDa. At pH ≤ 6.6, all three forms were mainly present in the cytoplasm and only trace amounts were observed in the nucleus. However, as pH increased to 7.7, the majority of the three isoforms localized to the nucleus (Fig 5A). Full-length PacC has two predicted proteolytic sites positioned at the 80th and 222th amino acid residue, respectively, (http://www.expasy.org/tools/peptidecutter/) (Fig 5B; S3 Table) that may lead to the production of 36 kDa and 52 kDa eGFP-fused peptides. To investigate the functions of these predicted truncated versions of PacC, two constructs, *eGFP-PacC^96^* (artificial truncate at the 96th amino acid) and *eGFP-PacC^245^(artificial* truncate at the 245th amino acid), were expressed under control of the native PacC promoter in a Δ*pacC* mutant to create transformants NGP96 and NGP245. The NGP96 transformants expressed a 36 kDa protein with a protein >36kDa. In the NGP245 transformant, a 52 kDa protein, and a 36 kDa protein were detected together with a protein > 52kDa. These data suggested that the 36 kDa and 52 kDa proteins may be generated from the full length eGFP-PacC^559^ by processing, probably at the 80th and 222th aa sites (Fig 5C), respectively.

To determine which PacC isoform is responsible for transcriptional activation or repression, three constructs were created in which the binding domain (BD) of the Gal4 protein on pGBKT7 was fused with the full-length PacC (*pBD-PacC^559^*), the truncated PacC^245^ (*pBD-PacC^245^*) and PacC^96^ (*pBD-PacC^96^*). Transformants expressing pBD-PacC^559^ or pBD-PacC^96^ were prototrophic for Trp but failed to grow on SD-Trp-His plates. In contrast, transformants of pBD-PacC^245^ grew well on the plates with galactosidase activity (Fig 5D), suggesting that PacC^245^ can produce a protein with transcription activation capability. We also expressed and purified GST fusion proteins of PacC^559^, PacC^222^ and PacC^80^ and verified that GST-PacC^559^ and GST-PacC^222^, but not GST-PacC^80^ could bind to the GCCAAG consensus sequence (Fig. 5E). These data suggest that PacC^222^ functions as a transcriptional activator, but PacC^559^ may act as a transcriptional suppressor, and PacC^80^ by itself probably does not act as a transcription factor.

To confirm the *in vivo* regulatory functions of PacC^559^ and PacC^222^, 22 genes with the PacC-binding GCCAAG consensus in their promoters were randomly selected for qRT-PCR analysis with total RNA from the WT, the Δ*pacC*, NGP245, and NGP559 strains. Based on their expression patterns, these genes were classified into three types (Fig 6). There were six type I genes, which were repressed under alkaline conditions in the WT and NGP559, but up-regulated in the Δ*pacC* mutant and NGP245 (Fig 6A), indicating that they are repressed by PacC^559^. Fourteen type II genes were up-regulated in the wild type, NGP245 and NGP559 strains, but significantly reduced in the Δ*pacC* mutant (Fig 6B), confirming that PacC^222^ is a transcription activator. We further assayed the phenotypes of NGP96, NGP245 and NGP559 strains. Sensitivity to alkaline pH, conidiation and virulence were fully complemented in NGP559, but only partially so in NGP245. No obvious difference was observed between the Δ*pacC* mutant and NGP96 (Fig 5F-5H). When considered together, these results provide evidence that PacC^222^ functions as a transcription activator, PacC^559^ as a transcription repressor, and that both activities are necessary for the full biological function of PacC during plant infection by *M. oryzae*.

**Fig 6.**
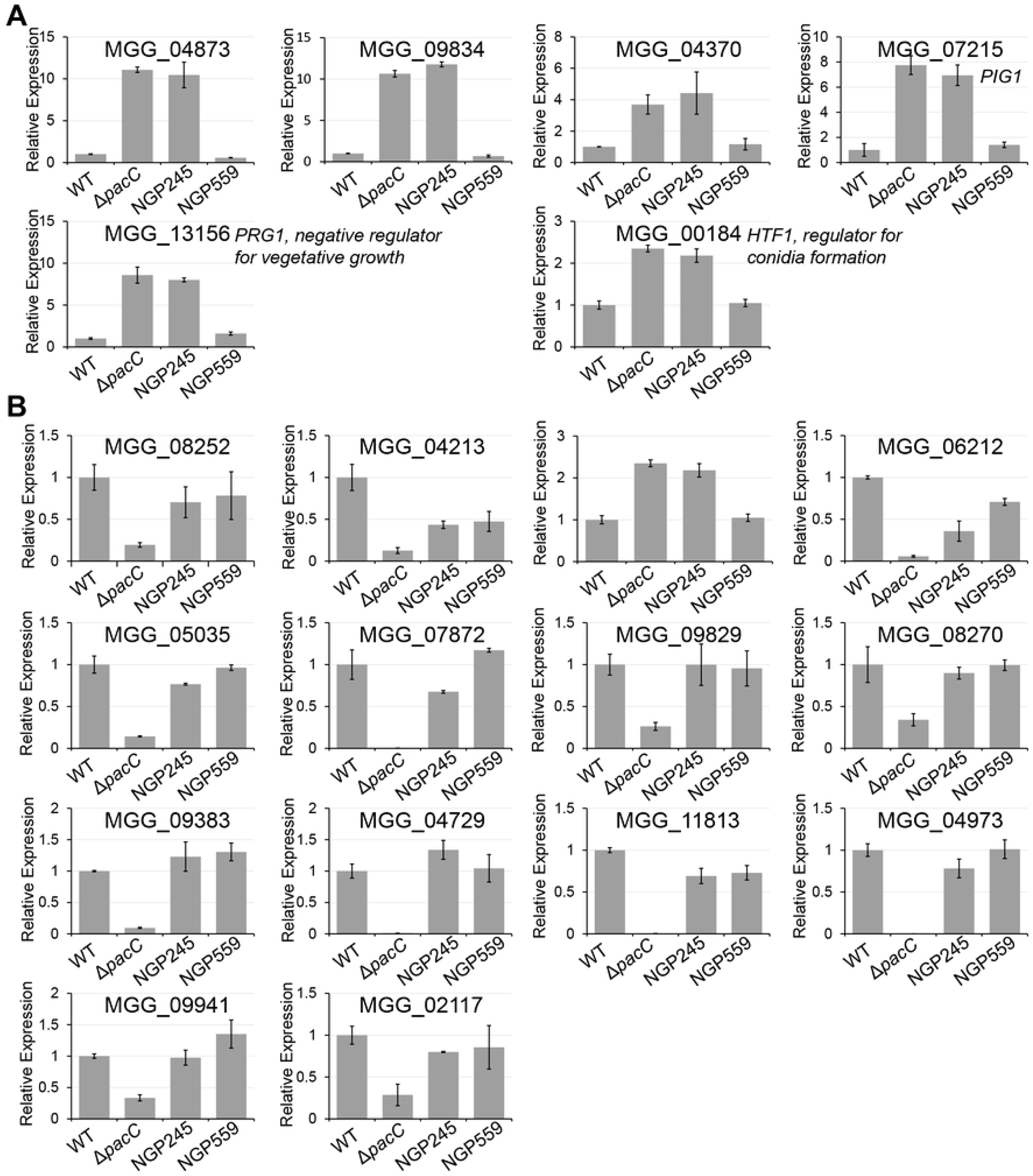
Bar charts showing qRT-PCR analysis to show genes down-regulated by PacC^559^ or up-regulated by PacC^222^. (A) Bar charts showing qRT-PCR analysis of genes up-regulated in the Δ*pacC* mutant and NGP245, but down-regulated in the wild type and NGP559. (B) Genes down-regulated in the Δ*pacC* mutant, but up-regulated in the wild type, NGP245 and NGP559. The relative expression levels of each gene were assayed by qRT-PCR with total RNA isolated from mycelium cultured under pH 7.7. For each gene, its expression level in the wild type P131 cultured in pH 7.7 was arbitrarily set as 1. WT, the wild-type P131; Δ*pacC, PacC* deletion mutant; NGP245, a transformant of the Δ*pacC* mutant with *eGFP-PacC^245^* construct; NGP559, a transformant of the Δ*pacC* mutant with *eGFP-PacC^559^* construct.

### PacC orchestrates distinct developmental processes by regulating a broad repertoire of transcription factor-encoding genes

To understand how *PacC* in *M. oryzae* affects the observed phenotypes, we analyzed the downstream hierarchy of transcriptional regulation. To do this, we selected several transcription factor-encoding genes directly regulated by PacC for functional analysis.

*MGG_01779* is a PacC-regulated Type II gene probably encoding a novel C6 zinc DNA binding domain protein (Fig 7), named *PAG*1 (for <u>P</u>acC <u>A</u>ctivated <u>G</u>ene 1). Its promoter has two GCCAAG sites bound by PacC (Fig 7B). The Δ*pag*1 mutant showed no obvious defects in colony growth or conidiation (Fig 7C and D, S4J Fig), but was significantly impaired in IH branching and in virulence (Fig 7E). *PAG*1 therefore functions downstream of PacC to regulate biotrophic growth of IH.

**Fig 7.**
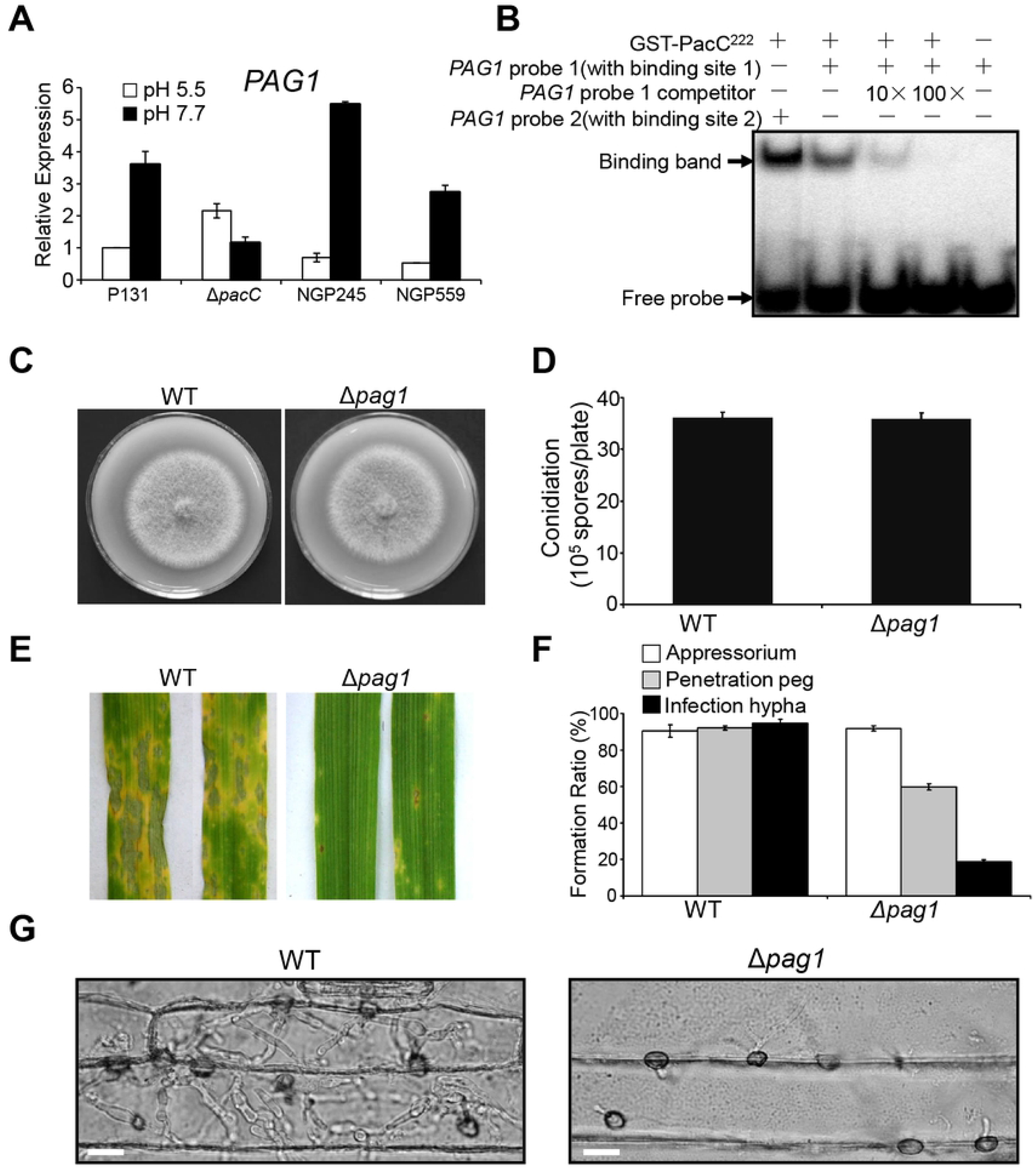
*PAG1* is positively regulated by PacC and important for biotrophic growth of *M. oryzae*. (A) Expression of *PAG1* in mycelium of WT P131, Δ*pacC*, NGP239, and NGP559. For the Q-PCR, the expression level of P131 cultured at pH 5.5 was arbitrarily set to 1. (B) PacC binds to GCCAAG motifs in the *PAG1* promoter. Purified GST-PacC^222^ protein was used to detect binding of putative PacC binding motifs. Probes 1 and 2 contain predicted PacC-binding motifs and were prepared by labelling with ^32^P-dCTP and incubated with GST-PacC^222^ for 30 min, before loading a native-PAGE gel. For competition experiments, 100x or 10x concentrations of un-labelled Probe 1 were mixed with the GST-PacC^222^ protein for 30 min before incubation with ^32^P-dCTP-labelled probes. (C) WT and Δ*pag1* are same in colony growth on OTA plates. (D) WT and Δ*pag1* are equivalent in conidiation. (E) Reduced virulence of Δ*pag1* mutant compared to WT. (F) Infection assays of WT and Δ*pag1* on barley epidermis. (G) Arrested biotrophic growth of Δ*pag1* compared to WT. Bar = 25 μm.

*MGG_13156* is a Type I gene (named *PRG1* for <u>P</u>acC <u>R</u>epressed <u>G</u>ene 1) with a GCCAAG site in its promoter (S5B Fig). It appears to be expressed at higher levels under acidic rather than alkaline conditions, and was up-regulated in the Δ*pacC* mutant (S5A Fig). We also generated two Δ*prg1* mutants, neither of which had obvious growth defects (S4I Fig). However, transformants showing de-repression of *PRG1* by PacC, produced smaller colonies (S5C-S5G Fig). Thus, *PRG1* is indeed negatively regulated by PacC during mycelial growth and infection.

*HTF1* is a homeobox gene required for conidiophore formation [51]. It was repressed in the WT, but up-regulated in the Δ*pacC* mutant under alkaline pH conditions and during plant infection (S6A Fig). Its promoter contains a GCCAAG site (S6B Fig). The *PIG1* TF controls mycelial melanin biosynthesis [52] and was repressed by PacC^559^ under alkaline pH (S7A Fig). Its promoter also has a GCCAAG site (S7B Fig), indicating that PacC represses expression of *PIG1* and thus the Δ*pacC* mutant formed darker colonies for excess melanization (Fig 2A).

Taken together, these results indicate that PacC may orchestrate distinct developmental processes in *M. oryzae*, by directly regulating a wide range of transcription factors, thereby leading to large-scale transcriptional changes which are essential for establishing blast disease.

## Discussion

Many of the most important plant pathogenic fungi are hemibiotrophs which are able to switch from biotrophic growth and necrotrophic growth during plant infection [1–3, 9–10,]. Very little, however, is known regarding how these fungi switch between the two growth habits and, indeed, which signals from the plant elicit such a dramatic developmental and physiological change. In this study, we identified host cellular pH as an inductive signal for the necrotrophic switch in *M. oryzae*.

We found that that host cells around infection sites are initially and temporally alkalinized to pH 7.8 during biotrophic growth, but acidification to pH 6.5 then follows during necrotrophic growth (Figure 1A-1B and 1D-1E). This observation contrasts with a previous study which indicated that host cells infected by *M. oryzae* remains alkaline for up to 60 hpi [51]. Our results also showed that pH alkalinization is independent of *PacC* function (Fig 1B-1C and 1E) and is clearly inhibitory to fungal growth and conidiation, based on *in vitro* studies (Fig 1F-1G), suggesting that host cellular alkalinization is a plant immune response to *M. oryzae*, as previously reported in other disease systems [18–21]. However, later acidification of plant tissue appears to be an active process, manipulated by *M. oryzae*, because a Δ*pacC* mutant of the fungus is deficient in its induction (Fig 1B-1C). Moreover, acidified pH is conducive to fungal growth and conidiation in vitro (Fig 1F-1G). Therefore, biotrophic growth may require a mechanism of fungal adaptation to host pH alkalinization while necrotrophic growth is an active process by which the fungus prepares for future propagation.

Many previous studies have showed that the PacC transcription factor is important for virulence of plant fungal pathogens [54–58], but our study, however, offers a potential mechanism which indicates that the PacC pathway (including the PacC transcription factor) is involved in the regulation of fungal hemibiotrophy. We have shown that the PacC pathway is required for *M. oryzae* not only to adapt to the host cellular alkalinized pH for biotrophic growth, but also to induce pH acidification, which allows necrotrophic growth (Fig 1; Fig 2). In particular, the regulation of hemibiotrophy involves shuttling of the PacC transcription factor between the nucleus and the cytoplasm, whereby PacC localizes to the nucleus during biotrophic growth and alkaline pH, but to the cytoplasm during necrotrophic growth and acidic conditions (Fig 2; Fig 5A). Interestingly, palH in *A. nidulans* mechanistically resembles mammalian GPCRs [59], and this study has revealed its requirement for virulence of *M. oryzae* (Fig. 2A-2E). When considered together, PalH there may be a candidate target for development of antifungal drugs.

This study also showed that *M. oryzae* differs from other fungi in the composition of the PacC pathway, and in the proteolytic processing of the PacC transcription factor. In *A. nidulans*, palI enhances the plasma membrane localization of palH [28], and palA and Vps32 form a complex with palB for proteolytic processing of pacC [32–33]. However, *M. oryzae* mutants lacking the three orthologous proteins are indistinguishable from the wild type strain in alkaline pH sensitivity, colony growth, conidiation and virulence (Fig.2), indicating that they are not required for PacC pH-signaling in *M. oryzae*. The PacC transcription factor in *M. oryzae* has two functional forms, a full-length transcriptional repressor and a medium truncated form that acts as transcription activator, both of which are required for its biological function (Fig 5 and Fig 6). To achieve their distinct transcriptional functions, *M. oryzae* PacC forms translocate to the nucleus during biotrophic growth (Fig 3A), likely as a consequence of alkaline pH (Fig 1A-1B; Fig 3B), as reported for nuclear localization of pacC in *A. nidulans* [37]. However, the two different *M. oryzae* PacC forms exist independently of alkaline pH (Fig 5A). Therefor the processing of PacC in *M. oryzae* is different from that reported in other fungi. In *A. nidulans*, pacC remains in the cytoplasm in its full-length closed conformation, which is protease-inaccessible under acidic conditions, but is processed into the activator form under alkaline pH (34-35,49). Furthermore, in *C. albicans*, the full length 85kDa CaRim101p is partially processed into a 74 kDa, under protein neutral pH, but into a 64kDa protein in acidic pH [42]; In *S. cerevisae*, the full length 98 kDa Rim101p is cleaved into an active 90kDa form in alkaline pH [60–61]. It is noteworthy that PalA and Vps32, which are involved together with PalB in pH-dependent proteolytic processing of PacC in other fungi, are dispensable in *M. oryzae* for both alkaline pH adaptation and virulence(Fig.2), suggesting that they are not essential for generating the different transcriptional regulator forms of PacC in *M. oryzae*. Whether the dispensability of PalA and Vps32 is related with acidic processing of PacC in *M. oryzae* needs to be further addressed.

This study has revealed that full length PacC in *M. oryzae* also acts as a transcription repressor, this is similar to the full-length form of Rim101p in *S. cerevisiae* [41]. In *C. albicans*, CaRim101p also functions as both an activator and a repressor and has multiple forms although it is unclear which form acts as a transcription repressor [43–44]. In *A. nidulans* there also is a full-length form of pacC in the nucleus under alkaline pH [37], and it is reported that *A. nidulans* pacC can repress the acid-expressed *gabA* gene by binding to its promoter regions and preventing binding of the transcriptional activator *IntA* [62]. This suggests that in *A. nidulans* PacC can function as a transcription repressor. However, it remains less clear how the two PacC isoforms recognize their specific targets and achieve selectivity in transcriptional activation and repression.

Our results provide evidence that *M. oryzae* PacC activates genes that are specifically expressed during biotrophic growth, and at the same time repress expression of genes that are related to saprophytic growth (Fig.4A-4C). Over 2700 genes are differentially expressed in biotrophic IH of *M. oryzae* when compared with a Δ*pacC* mutant (Fig. 4A; S1 Dataset). Therefore, more than 25% of the total protein-encoding genes in *M. oryzae* genome are regulated by PacC. This is much higher than the number regulated by PacC in other plant pathogenic fungi [54, 56], where mycelium grown in axenic conditions was used for RNA-Seq analysis. This is, however, the first time that biotrophic infection has been analyzed for PacC regulation. We confirmed that *M. oryzae* PacC can bind to the GCCAAG *cis*-element, as previously reported for *A. nidulans* PacC (Fig.5E, Fig.7B, S5B Fig., S6B Fig., S7B Fig) [38]. By surveying PacC binding motifs in promoters of each differentially expressed gene, we have identified nearly 1500 genes that are very likely to be directly regulated by PacC in *M. oryzae* biotrophic IH (Fig 4A; S4 Dataset). It is notable that among the putatively direct targets of PacC are genes enriched in those encoding extracellular and plasma membrane proteins (Fig 4D-4F; S4 Dataset), that are likely to be involved in suppression of plant immunity, remodeling fungal cell walls and acquisition of carbon and nitrogen sources from living host cells (Fig 4C-4D; S4 Dataset; S5 Dataset) [50]. In particular, more than 80 previously characterized virulence genes appear to be regulated by PacC (S6 Dataset), including effector protein genes and 12 genes that have been identified to participate in acquisition and utilization of nutrients. Taken together, it seems likely that an important role of PacC in *M. oryzae* is to regulate expression of genes that suppress plant immunity, remodel the cell wall of invasive hyphae and facilitate acquisition and utilization of nutrients to allow biotrophic growth in alkalinized host plant cells.

In conclusion, we propose a model for the regulation of hemibiotrophic growth of *M. oryzae* by the PacC pH signaling pathway, as shown in Figure 8. Initially, *M. oryzae* penetration leads to pH alkalinization in host cells around infection sites (Fig 8A). The alkalinized host environment is sensed by Pal components of the PacC pathway, resulting in localization of PacC to the nucleus where the activator isoform enhances gene expression associated with biotrophic growth, while the repressor isoform represses genes associated with necrotrophic growth and conidiation (Fig 8B). After extensive tissue colonization, *M. oryzae* induces disintegration and acidification of host plant cells, which induces the two PacC isoforms to translocate from the nucleus thereby de-repressing gene expression associated with necrotrophic growth and conidiation (Fig 8C). The shuttling of PacC between the nucleus and cytoplasm according to pH in host tissue may thereby regulate the temporal switch between biotrophic and necrotrophic growth that is essential for blast disease.

**Fig 8.**
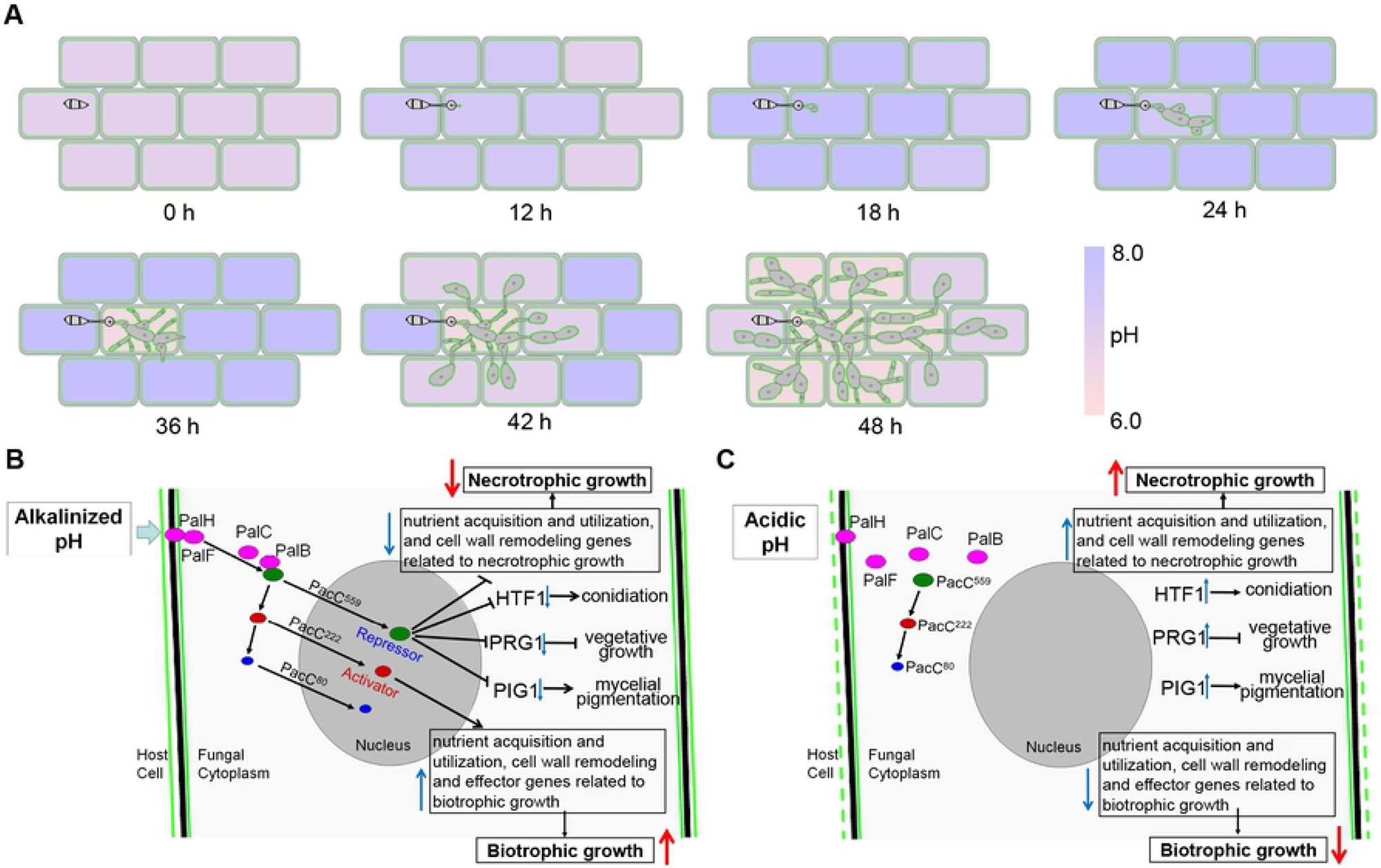
A Model for PacC-Dependent Regulation of Gene Expression Associated with the Biotrophic/Necrotrophic Switch during Infection of *M. oryzae*. (A) Infected plant cells are alkalinized during the early biotrophic growth of *M. oryzae* and then become acidified during the later necrotrophic growth. (B) During biotrophic growth, the PacC^559^ and PacC^222^ transcription factor isoforms localize to the nucleus, where PacC^559^ acts as a transcriptional repressor to repress expression of genes associated with conidiation and necrotrophic growth, including *PRG1*, *HTF1*, and *PIG1* while PacC^222^ acts as a transcriptional activator to activate genes associated with biotrophic growth. (C) As host cells become acidified and lose viability, the PacC functional isoforms exit from the nucleus thereby de-repressing expression of genes related to necrotrophic growth and conidiation.

## Materials and Methods

### Strains and culture conditions

The P131 strain of *M. oryzae* was used for all genetic manipulation and infection assays [63]. S1528, which has the opposite mating type to P131, was used only for genetic crossing and co-segregation analyses, as previously described [64]. Strains 70-15 and DG-ZX-83, together with P131, were only used to assay effect of pH on colony growth and conidiation. All the wild-type strains and transformants (Supplemental Table 4) were maintained on oatmeal tomato agar (OTA) plates at 28°C, as described [63]. For assaying colony growth under normal condition, mycelial blocks (φ=5 mm) were placed in the centre of complete medium (CM) plates, and cultured for 120 h at 28°C. Conidia were produced on OTA, as previously reported [65]. For assaying the effect of pH on colony growth and conidiation, CM plates and oatmeal agar plates were used, respectively, which were adjusted to different pH with appropriate buffers, as described in Supplemental Figure 2.

### Plant infection and microscopy observations

Rice and barley seedlings were grown, inoculated and incubated, as described for assaying virulence [14, 65]. Lesions on rice and barley leaves were examined at five days and three days post inoculation (dpi), respectively. To assay virulence on wounded rice leaves, detached rice leaves were scratched against leaf veins with a needle, then mycelial blocks (φ=2 mm) were placed onto wounded sites, and incubated in a moist chamber for 3 days.

Assays of the barley epidermal infection process, host-derived ROS generation and growth of infection hyphae, were performed as described by Chen et al. [14]. To investigate subcellular localization of PacC, microscopy was performed with a Δ*pacC/GFP:PacC* strain cultured in CM at different pH, or inoculated onto barley epidermis. To visualize the viability of barley epidermal cells at infection sites, Trypan blue staining of barley leaves was performed, as described previously [46]. All microscopic observations were made with a Nikon 90i epifluorescence microscope (Nikon, Japan).

### Molecular manipulations with DNA and RNA

Fungal genomic DNA was extracted, as described previously [66]. Total RNA was extracted with the TRIzol Plus RNA Purification Kit (Life technologies, USA). Standard molecular procedures were followed for plasmid preparation, Southern genomic DNA hybridization, and enzymatic digestion of DNA [67]. TAIL-PCR was performed as described according to Liu et al. [48]. The qRT-PCR reactions were performed with a ABI PRISM 7900HT system (Applied Biosystems, CA, USA) using the SYBR^®^ PrimeScript^™^ RT-PCR Kit (Takara, Dalian, China).

### Generation of gene deletion mutants and transformation

Fungal protoplasts were isolated from *M.oryzae* mycelium that was cultured in liquid CM at 180 rpm for 36 h and transformed as described [68]. CM plates supplemented with hygromycin B at 250 μg/ml (Roche, USA) or neomycin at 400 μg/ml (Amresco, USA), were used for selecting hygromycin-resistant or neomycin-resistant transformants.

Mutants with targeted deletions of *PalI*, *PalA*, *PalB*, *PalC*, *PalF*, *PalH*, *PacC* and Vps32 were generated by one-step targeted gene replacement. To construct a gene replacement vector for each gene, 1.5 kb upstream and 1.5 kb downstream sequences were amplified and cloned into pKNH [63]. The resulting vectors were independently transformed into P131. Transformants resistant to hygromycin, but sensitive to neomycin, were subjected to screening by PCR. Gene deletion candidates were further confirmed by Southern blot analysis.

For genetic complementation, genomic DNA fragments of individual *PalB*, *PalC*, *PalF*, *PalH*, and *PacC* genes containing 1.5 kb promoter and 0.5 kb terminator regions were amplified and cloned into pKN [63]. To generate the *eGFP-PacC* fusion construct pKGPacC^559^, a fragment amplified with primers PacCNP5 and PacCNP3 was digested with *Xho*I and *Hin*dIII and cloned into pKNTG [63]. The same strategy was used to generate the pKGPacC^245^ and pKGPacC^96^ constructs. pKGPacC^559^, pKGPacC^245^ and pKGPacC^96^ were independently transformed into a Δ*pacC* mutant to generate strains NGP559 (Δ*pacC/GFP:PacC*), NGP245 and NGP96, respectively. pKGPacC^559^ was also transformed into Δ*palH, ΔpalF, ΔpalC* and Δ*palB* to generate ΔpalH/*GFP:PacC*, ΔpalF/*GFP:PacC*, ΔpalC/*GFP:PacC* and Δ*palB/GFP:PacC* strains, respectively.

### RNA-seq analysis

Total RNAs were isolated with the TRIzol Plus RNA Purification Kit (Life technologies, USA) from epidermis of barley leaves that were inoculated with the P131 and the Δ*pacC* strain at 18 hpi. Each RNA sample was subjected to DNase digestion (TaKaRa, Dalian, China), to remove DNA contamination and then mRNA was purified with poly-T oligo-attached magnetic beads. Construction of the libraries and RNA-Seq analysis were performed by Novogene Corporation (Beijing, China) using an Illumina HiSeq 2000 platform (Illumina, Inc., USA). Raw reads were filtered to remove adaptor and low-quality sequences. Clean reads were mapped to the P131 genome [69] using Tophat2 [70], allowing up to two base mismatches. The <u>F</u>ragments <u>P</u>er <u>K</u>ilobase of transcript sequence per <u>M</u>illions base pairs sequenced (FPKM) was used to indicate expression levels of *M. oryzae* genes, which were calculated with Cuffdiff [71]. For either of the strains, two independent sets of inoculated epidermis were prepared to generate the RNA-seq data. The expression levels of *MGG_03982* (Actin gene) were used for normalizing different samples. Differentially expressed genes between the wild type and the Δ*pacC* mutant were defined under the criteria of the absolute log2 fold change value ⩾ 0.5, q value ⩽ 0.05 and FPKM ⩾ 2 in at least one of the strains.

### Intracellular pH measurement of barley leaf epidermal cells

The lower epidermis of barley leaves, inoculated with conidial suspension, was removed at designated times to measure ambient pH in infected cells using 2’,7’-bis-(2-carboxyethyl)-5-(and-6)-carboxyfluorescein-acetoxymethyl ester (BCECF-AM) fluorescein as a pH indicator dye [45]. Emission intensities at 530 nm and 640 nm of the dye after excitation at 488 nm were used to calculate pH, in accordance with a calibration curve between the fluorescence ratio (F530/F640) and pH change, which was established with healthy barley leaf epidermis, as described in the supplemental data. Three independent experiments were performed. For each of the experiments, pH was measured in at least ten single plant cells underneath conidia or appressoria or with invasive hyphae.

### Transcription activation assays

To assay transcriptional activation activities of PacC, the full-length cDNA of *PacC* amplified with primers PacC-F and PacC-R was digested with *Eco*RI and cloned into pGBKT7 (Clontech, USA) as pBD-PacC^559^. A fragment encoding PacC^245^ and PacC^96^ were amplified with primers PacC-F/PacCR245 and PacC-F/PacCR96, and cloned into vector pGBKT7 as pBD-PacC^245^ and pBD-PacC^96^, respectively. These vectors were individually transformed into the yeast strain AH109. Yeast cells were grown on SD-Trp and SD-Trp-His plates. Transformants with the empty vectors pGBKT7 and pGBT9 were used as the negative and positive control, respectively. After 3 days incubation at 28°C, X-α-gal was added to SD-Trp-His plates for assaying β-galactosidase activities.

### Electrophoretic mobility shift assays (EMSA)

GST-fused PacC^559^, PacC^222^ and PacC^80^ proteins were individually expressed in the pGEX-4T-3 vector (Promega, USA) in *E. coli* BL21DE3 and purified, as described in supplementary data. Double-stranded oligonucleotides were used as DNA templates for EMSA, which were formed by mixing equal amounts of two single complementary oligonucleotides in the annealing buffer (0.2 M Tris-HCl, 1 mM EDTA, 0.5 M NaCl, pH 8.0), heating for 5 min at 95 °C, and cooling down to room temperature. The double-stranded oligonucleotides were end-labeled with [α-^32^P]-dCTP with the Random Primer Labeling Kit (Takara, Dalian, China). The binding reaction (28 μl) was performed in binding buffer (10 mM Tris-HCl, pH 7.5, 5 mM NaCl, 1 mM DTT, 1 mM EDTA, 5% glycerol) with 50 ng of purified GST-fused PacC proteins and 2 pM labeled template DNA. 20 pM and 200 pM un-labelled DNAs were used as 10× and 100× specific competitors, respectively. The samples were separated on 8% native PAGE gels for 50 min, which were exposed to X-ray film for 1 h and detected by a storage phosphor system (Cyclone, PerkinElmer, USA).

### Isolation of cytoplasmic and nuclear proteins from mycelia of *M. oryzae*

Cytoplasmic and nuclear proteins were isolated from protoplasts, as described with some modifications [72]. The protoplasts were suspended in 400 μl buffer A (10 mM HEPES-KOH, pH 7.9, 1.5 mM MgCl_2_, 10 mM KCl, 0.5 mM DTT, 0.2 mM PMSF, 1 mg/ml leupeptin, 1 mg/ml pepstatin) and vortexed six times. Each vortex lasted for 2 minutes, and 3 minute intervals were allowed between each vortex to incubate the suspension on ice. After vortexing, the mixture was centrifuged at 4,000 g for 15 min at 4°C to collect the supernatant as cytoplasmic proteins (CP). Then 200 μl buffer B (20 mM HEPES-KOH, pH 7.9, 25% glycerol, 1.5 mM MgCl_2_, 420 mM NaCl, 0.2 mM EDTA, 0.5 mM DTT, 0.2 mM PMSF, 1 mg/ml leupeptin, 1 mg/ml pepstatin) was added to suspend the pellets on ice, which was vortexed again as mentioned above, and centrifuged at 12,000 g for 30 min at 4°C to collect the supernatant as nuclear proteins (NP). The CP and the NP were mixed in ratio of 2:1 as the total proteins (TP). 30 μl TP, 20 μl CP, and 10 μl NP were mixed with equal volume 2×loading buffer and separated on 10% SDS-PAGE. Western blot was performed with an anti-GFP antibody (Abmart, China).

### Plasmid Construction

Cloning strategies for all plasmid constructions are described in the Extended Experimental Procedures. All primers used and plasmids constructed were listed in Supplemental Tables 5 and 6, respectively.

### Accession numbers

Sequence data from this study can be found in the GenBank/EMBL data libraries under accession number HQ889837 (*PacC*), HQ889838 (*PalH*), HQ889839 (*PalF*), HQ889840 (*PalC*), and HQ889841 (*PalB*).

## Acknowledgments

This work was supported by grants from the Ministry of Science and Technology (2016YFD0300700, 2012CB114000), the Ministry of Agriculture and Rural Affairs (CARS-01-16, 201203014) and the Ministry of Education (B13006, IRT1042), China, to Y.-L.P. We thank Professor Xinnian Dong at Duke University for her critical reading of the manuscript.

## Supporting Information

**S1 Fig. *In situ* pH calibration curve and three-dimension visualization of cellular pH in barley leaf epidermal cells infected with *M. oryzae*.**

(A) A calibration curve of the fluorescence ratio (F530/F640) against pH changes was established by using epidermis of un-inoculated barley leaves. The epidermis was pre-treated for 15 min with Nigericin (Invitrogen, USA) dissolved in buffers (pH 6.0, 6.5, 7.0, 7.5 and 8.0) and then added with the dye, BCECF-AM fluorescein, before visualization by laser scanning confocal microscopy. Emission intensities at 530 nm and 640 nm of the dye after excitation at 488 nm, were recorded with a Nikon A1 Laser scanning confocal microscope (Nikon, Japan). Error bars represent the standard deviation of 14 images taken from 7 different epidermal cells. (B) Three-dimension visualization of cellular pH in barley leaf epidermal cells infected by *M. oryzae*. A bright-field and Ratiometric CSLM image of BCECF-AM stained barley epidermal cells infected by the wild type P131 at 18 hpi (up panel) and at 36 hpi (bottom panel). Cross sections were also made, as indicated in the red and blue lines and are shown at the right and bottom, respectively. PIH, primary infection hyphae; BIH, branched infection hyphae; CSLM: confocal scanning laser microscopy; Bar = 25 μm. BCECF-AM: 2’,7’-bis-(2-carboxyethyl)-5-(and-6)-carboxyfluorescein-acetoxymethyl ester.

**S2 Fig. Effect of pH on colony growth and conidiation of *M. oryzae* isolates 70-15 and DG-ZX-83.** Colony growth and conidiation by *M. oryzae* were assayed on complete medium (CM) and oat broth (OBA) medium, respectively. OBA medium was made from agar (16g/L) and oat broth, which was prepared by boiling flattened oats (10 g/L) in water for 20min and filtering with gauze. CM and OBA were adjusted to pH 5.5 with 0.2M sodium acetate buffer (final concentration 25mM), to pH 6.0 - 7.5 with 0.2M phosphate buffer solution (final PO_4_^3-^concentration 50mM), or to pH 7.8 to 8.5 with 0.2M Tris-HCl buffer (final concentration to 50mM). (A) Graphs showing colony growth of 70-15 and DG-ZX-83 under different pH conditions. (B) Conidiation of 70-15 and DG-ZX-83 under different pH conditions. (C) Colony morphology of 70-15 and DG-ZX-83 under different pH conditions. (D) Micrographs of conidium formation of 70-15 and DG-ZX-83 under different pH conditions. Colony diameter and conidium number formed under individual pH conditions were normalized against those formed at pH 6.5. Bar = 20 μm.

**S3 Fig. Phenotypes of nine *Agrobacterium tumefaciens-mediated* transformation (ATMT) mutants of *M. oryzae* and positions of their T-DNA insertion sites.** (A) Colony growth of ATMT mutants compared with the wild type P131 on oatmeal tomato agar (OTA) plate and CM plates buffered at pH 6.6 and 7.7. Colonies were photographed at 120 hpi. (B) Conidia production of ATMT mutants as compared with the wild type P131. Conidia were harvested from the strains that were cultured on OTA plates (Φ=6 cm). Means and standard deviation were calculated from three independent experiments. \*\**p < 0.01, n* > 100. (C) Virulence of ATMT mutants compared to the wild type P131. Conidia of P131 and the mutants with concentration of 5×10^4^ spores/ml in 0.025% Tween 20 were used to spray the barley leaves. Infected leaves were photographed at 5 dpi. (D) Diagram showing T-DNA integration sites in the nine alkaline pH-sensitive ATMT mutants. The insertion sites are marked with black arrows and numbers indicating the relative distance to the ATG codon of corresponding genes.

**S4 Fig. Strategies for generating gene deletion mutants of the PacC pathway and corresponding Southern blot analysis.** To generate gene replacement constructs, approximately 1.5 kb of upstream and 1.5 kb of downstream flanking sequences (shaded in gray) of each targeted gene were amplified with specific primer pairs listed in Table S9. The resulting PCR products were cloned into restriction enzyme sites flanking the hygromycin phosphotransferase (*hph*) gene of plasmid pKNH to generate specific gene replacement vectors (left panes of A-G). The right panels in (A)-(G) are images of Southern blots of the resulting knockout mutants hybridized with the probes marked in the schematic drawings. Genomic DNAs were isolated from the wild-type strain P131 (WT) and two or three representative knockout mutants for each gene. B, *Bam*HI; C, *Cla*I; E, *Eco*RI; H, *Hin*dIII; K, *KpnI;* P, *PstI;* S, *SpeI;* SA, *SacI;* X, *XhoI*.

**S5 Fig. *PRG1* is negatively regulated by PacC and its overexpression results in smaller colony growth.** (A) Expression of *PRG1* in mycelia of WT P131, Δ*pacC*, NGP239, and NGP559. (B) PacC binding to the GCCAAG consensus from the promoter of *PRG1*. (C) Colonies of P131 (WT), the Δ*prg1* mutant, RP8 and MC1. RP8 and MC1 are a *PRG1* overexpression transformant driven by the RP27 promoter and a transformant expressing the mutant allele of *PRG1* with the PacC-binding site changed from GCCAAG to CTGCAG in its native promoter of the Δ*prg1* mutant, respectively. (D) Colony diameter of following strains: WT, the Δ*prg1* mutant, transformants RP2, RP8, RP12, RP14, RP15, RP18, RP20 that overexpress *PRG1* driven by the RP27 promoter, transformants MC1, MC6, MC11, MC19, MC20, MC21, MC28 that over-express *PRG1* driven by its native promoter with the PacC-binding cis-element mutated as described in (C). (E) Expression levels of *PRG1* in the same set of strains described above. (F) Conidiation of WT and the Δ*prg1* mutant. (G) Virulence of WT and the Δ*prg1* mutant on barley leaves.

**S6 Fig. *HTF1* is negatively regulated by PacC559.** (A) Quantitative RT-PCR analysis of *HTF1* expressed in mycelia of different strains under alkaline and acidic pH. (B) PacC binding to the GCCAAG consensus from the promoter of *HTF1*.

**S7 Fig. *PIG1* is negatively regulated by PacC.** (A) Quantitative RT-PCR analysis of *PIG1* expressed in mycelia of different strains under alkaline and acidic pH. (B) PacC binding to the GCCAAG consensus from the promoter of *PIG1*.

**S1 Table.** Co-segregation of the phenotype changes with the hygromycin resistance in the nine alkaline pH-sensitive mutants.

**S2 Table.** Colony sizes of the WT strain and deletion mutants grown at different ambient pH.

**S3 Table.** Putative PacC cleavage sites predicted by the Peptidecutter.

**S4 Table.** *M. oryzae* Strains used in this study.

**S5 Table.** Plasmids used in this study.

**S6 Table.** Primers used in this study.

**S1 Dataset.** Genes expressed in biotrophic infection hyphae of the wild type and Δ*pacC* strains at 18hpi.

**S2 Dataset.** Expression levels in other stages of the PacC-activated genes during biotrophic growth.

**S3 Dataset.** Expression levels in other stages of the PacC directly suppressed genes during biotrophic growth.

**S4 Dataset.** Annotations of genes directly regulated by PacC in the biotrophic infection hyphae.

**S5 Dataset.** Glycoside hydrolases may act on plant cell wall polysaccharides to provide carbon sources for biological growth of infection hyphae

**S6 Dataset.** Previously reported pathogenicity-important genes that are differentially regulated by PacC in the biotrophic infection hyphae.

## References

1. Oliver, R.P., and Ipcho, S.V. (2004). *Arabidopsis* pathology breathes new life into the necrotrophs-vs.-biotrophs classification of fungal pathogens. Mol. Plant. Pathol. 5, 347–352.

2. Rowe, H.C., and Kliebenstein, D.J. (2010). All mold is not alike: the importance of intraspecific diversity in necrotrophic plant pathogens. PLoS Pathog. 6, e1000759.

3. Spanu, P.D. (2012). The genomics of obligate (and nonobligate) biotrophs. Ann. Rev. Phytopathol. 50, 91–109.

4. Fernandez, J., Marroquin-Guzman, M. and Wilson R.A. (2014) Mechanisms of nutrient acquisition and utrilization during fungal infections of leaves. Annu. Rev. Phytopathol. 52:155–74

5. Glazebrook, J. (2005). Contrasting mechanisms of defense against biotrophic and necrotrophic pathogens. Annu. Rev. Phytopathol. 43, 205–227.

6. Sun, G., Elowsky, C., Li, G. and Wilson, R.A. (2018) TOR-autophagy branch signaling via Imp1 dictates plant-microbe biotrophic interface longevity. PLoS Genetics 14: e1007814

7. Dean, R., Van Kan, J.A., Pretorius, Z.A., Hammond-Kosack, K.E., Di Pietro, A., Spanu, P.D., Rudd, J.J., Dickman, M., Kahmann, R., Ellis, J., and Foster, G.D. (2012). The Top 10 fungal pathogens in molecular plant pathology. Mol. Plant Pathol. 13(4), 414–430.

8. Fernandez, J. and Orth, K. (2018) Rice of a Cereal Killer: The biology of *Magnaporthe oryzae* biotrophic growth. Trends Microbiol. 26, 582–97.

9. Kankanala, P., Czymmek, K., and Valent, B. (2007). Roles for rice membrane dynamics and plasmodesmata during biotrophic invasion by the blast fungus. Plant Cell 19, 706–724.

10. Wilson, R. A., and Talbot, N. J. (2009). Under pressure: investigating the biology of plant infection by *Magnaporthe oryzae*. Nat. Rev. Microbiol. 7, 185–195.

11. Giraldo, M.C., Dagdas, Y.F., Gupta, Y.K., Mentlak, T.A., Yi, M., Martinez-Rocha, A.L., Saitoh, H., Terauchi, R., Talbot, N.J., and Valent, B. (2013). Two distinct secretion systems facilitate tissue invasion by the rice blast fungus *Magnaporthe oryzae*. Nat. Commun. 4, 1996.

12. Khang, C.H., Berruyer, R., Giraldo, M.C., Kankanala, P., Park, S.Y., Czymmek, K., Kang, S., and Valent, B. (2010). Translocation of *Magnaporthe oryzae* effectors into rice cells and their subsequent cell-to-cell movement. Plant Cell 22, 1388–1403.

13. Mentlak, T.A., Kombrink, A., Shinya, T., Ryder, L.S., Otomo, I., Saitoh, H., Terauchi, R., Nishizawa, Y., Shibuya, N., Thomma, B.P., and Talbot, N.J. (2012). Effector-mediated suppression of chitin-triggered immunity by *Magnaporthe oryzae* is necessary for rice blast disease. Plant Cell 24, 322–335.

14. Chen, X.L., Shi, T., Yang, J., Shi, W., Gao, X.S., Chen, D., Xu, X.W., Xu, J.R., Talbot, N.J., and Peng, Y.L. (2014). N-glycosylation of effector proteins by an α-1,3-mannosyltransferase is required for the rice blast fungus to evade host innate immunity. Plant Cell 26(3), 1360–1376.

15. Sakulkoo, W., Oses-Ruiz, M., Oliveira Garcia, E., Soanes, D.M., Littlejohn, G.R., Hacker, C., Correia, A., Valent, B., and Talbot, N.J. (2018). A single fungal MAP kinase controls plant cell-to-cell invasion by the rice blast fungus. Science 359, 1399–1403.

16. Marroquin-Guzman, M., Hartline, D., Wright, J.D., Elowsky, C., Bourret, T.J., and Wilson, R.A. (2017). The *Magnaporthe oryzae* nitrooxidative stress response suppresses rice innate immunity during blast disease. Nat. Microbiol. 2, 17054.

17. Vylkova, S. (2017). Environmental pH modulation by pathogenic fungi as a strategy to conquer the host. PLoS Pathog. 13, e1006149.

18. Felix, G., Regenass, M., and Boller, T. (1993). Specific perception of subnanomolar concentrations of chitin fragments by tomato cells: induction of extracellular alkalinization, changes in protein phosphorylation, and establishment of a refractory state. Plant J. 4, 307–316.

19. Felle, H.H. (2001). pH: signal and messenger in plant cells. Plant Biol. 3, 577–591.

20. Zipfel, C., Kunze, G., Chinchilla, D., Caniard, A., Jones, J.D., Boller, T., and Felix G. (2006). Perception of the bacterial PAMP EF-Tu by the receptor EFR restricts *Agrobacterium*-mediated transformation. Cell 125, 749–760.

21. Gust, A. A., Biswas, R., Lenz, H.D., Rauhut, T., Ranf, S., Kemmerling, B., Götz, F., Glawischnig, E., Lee, J., Felix, G., and Nürnberger, T. (2007). Bacteria-derived peptidoglycans constitute pathogen-associated molecular patterns triggering innate immunity in *Arabidopsis*. J. Biol. Chem. 282, 32338–32348.

22. Fernandes, T.R., Segorbe, D., Prusky, D., and Di Pietro, A. (2017). How alkalinization drives fungal pathogenicity. PLoS Pathog. 13, e1006621.

23. Bateman, D.F., and Beer, S.V. (1965). Simultaneous production and synergistic action of oxalic acid and polygalacturonase during pathogenesis by *Sclerotium rolfsii*. Phytopathology 55, 204–211.

24. Cessna, S., Sears, V., Dickman, M., and Low, P. (2000). Oxalic acid, a pathogenicity factor of *Sclerotinia sclerotiorum,* suppresses the host oxidative burst. Plant Cell 12, 2191–2199.

25. Masachis, S., Segorbe, D., Turrà, D., Leon-Ruiz, M., Fürst, U., Ghalid, M.E., Leonard, G., López-Berges, M.S., Richards, T.A., Felix, G., and Pietro, A.D. (2016). A fungal pathogen secretes plant alkalinizing peptides to increase infection. Nature Microbiol. 1, 16043.

26. Peñalva, M.A., Tilburn, J., Bignell, E., and Arst, H.N., Jr. (2008). Ambient pH gene regulation in fungi: making connections. Trends Microbiol 16, 291–300.

27. Galindo, A., Calcagno-Pizarelli, A.M., Arst, H.N. Jr., and Peñalva, M.A. (2012) An ordered pathway for the assembly of ESCRT-containing fungal ambient pH signalling complexes at the plasma membrane. J Cell Sci. 125, 1784–1795.

28. Calcagno-Pizarelli, A.M., Negrete-Urtasun, S., Denison, S.H., Rudnicka, J.D., Bussink, H.J., Munera-Huertas, T., Stanton, L., Hervas-Aguilar, A., Espeso, E.A., Tilburn, J., Arst, H.N. Jr., and Peñalva M.A. (2007). Establishment of the ambient pH signaling complex in Aspergillus nidulans: PalI assists plasma membrane localization of PalH. Eukaryot Cell 6, 2365–2375.

29. Herranz, S., Rodriguez, J.M., Bussink, H.J., Sanchez-Ferrero, J.C., Arst, H.N., Jr., Peñalva, M.A., and Vincent, O. (2005). Arrestin-related proteins mediate pH signaling in fungi. Proc Natl Acad Sci U S A. 102, 12141–12146.

30. Hervas-Aguilar, A., Galindo, A., and Peñalva, M.A. (2010). Receptor-independent Ambient pH signaling by ubiquitin attachment to fungal arrestin-like PalF. J Biol Chem. 285(23), 18095–18102.

31. Galindo, A., Hervas-Aguilar, A., Rodriguez-Galan, O., Vincent, O., Arst, H.N., Jr., Tilburn, J., and Peñalva, M.A. (2007). PalC, one of two Bro1 domain proteins in the fungal pH signalling pathway, localizes to cortical structures and binds Vps32. Traffic 8, 1346–1364.

32. Denison, S.H., Orejas, M., and Arst, H.N., Jr. (1995). Signaling of ambient pH in Aspergillus involves a cysteine protease. J Biol Chem. 270, 28519–28522.

33. Peñas, M.M., Hervás-Aguilar, A., Munéra-Huertas, T., Reoyo, E., Peñalva, M.A., Arst, H.N., Jr, and Tilburn, J. (2007). Further characterization of the signaling proteolysis step in the *Aspergillus nidulans* pH signal transduction pathway. Eukaryot. Cell 6, 960–970.

34. Espeso, E.A., Roncal, T., Diez, E., Rainbow, L., Bignell, E., Alvaro, J., Suarez, T., Denison, S.H., Tilburn, J., Arst, H.N. Jr., and Peñalva, M.A. (2000). On how a transcription factor can avoid its proteolytic activation in the absence of signal transduction. EMBO J. 19, 719–728.

35. Diez, E., Alvaro, J., Espeso, E.A., Rainbow, L., Suarez, T., Tilburn, J., Arst, H.N., Jr., and Peñalva, M.A. (2002). Activation of the *Aspergillus* PacC zinc finger transcription factor requires two proteolytic steps. EMBO J. 21, 1350–1359.

36. Vincent, O., Rainbow, L., Tilburn, J., Arst, H.N., and Peñalva, M.A. (2003). YPXL/I is a protein interaction motif recognized by *Aspergillus* PalA and its human homologue, AIP1/Alix. Mol Cell Biol. 23, 1647–1655.

37. Mingot, J.M., Espeso, E.A., Diez, E., and Peñalva, M.A. (2001). Ambient pH signaling regulates nuclear localization of the *Aspergillus nidulans* PacC transcription factor. Mol Cell Biol. 21, 1688–1699.

38. Espeso, E.A., Tilburn, J., Sanchez-Pulido, L., Brown, C.V., Valencia, A., Arst, H.N. Jr, and Peñalva, M.A. (1997). Specific DNA recognition by the *Aspergillus nidulans* three zinc finger transcription factor PacC. J. Mol. Biol. 274, 466–480.

39. Fernandez-Martinez, J., Brown, C.V., Diez, E., Tilburn, J., Arst, H.N., Jr., Peñalva, M.A., and Espeso, E.A. (2003). Overlap of nuclear localisation signal and specific DNA-binding residues within the zinc finger domain of PacC. J Mol Biol. 334, 667–684.

40. Davis, D. (2003). Adaptation to environmental pH in *Candida albicans* and its relation to pathogenesis. Curr Genet. 44, 1–7.

41. Lamb, T. M., and Mitchell, A. P. (2003). The transcription factor Rim101p governs ion tolerance and cell differentiation by direct repression of the regulatory genes *NRG1* and *SMP1* in *Saccharomyces cerevisiae*. Mol. Cell Biol. 23, 677–686.

42. Li, M., Martin, S.J., Bruno, V.M., Mitchell, A.P., and Davis, D.A. (2004). *Candida albicans* Rim13p, a protease required for Rim101p processing at acidic and alkaline pHs. Eukaryot Cell. 3, 741–751.

43. Baek, Y.U., Martin, S.J., and Davis, D.A. (2006). Evidence for novel pH-dependent regulation of *Candida albicans* Rim101, a direct transcriptional repressor of the cell wall beta-glycosidase Phr2. Eukaryot. Cell 5, 1550–1559.

44. Ramon, A.M., and Fonzi, W.A. (2003). Diverged binding specificity of Rim101p, the *Candida albicans* ortholog of PacC. Eukaryot Cell. 2, 718–728.

45. Ozkan, P., and Mutharasan, R. (2002). A rapid method for measuring intracellular pH using BCECF-AM. Biochim. Biophys Acta 1572, 143–148.

46. Lipka, V., Dittgen, J., Bednarek, P., Bhat, R., Wiermer, M., Stein, M., Landtag, J., Brandt, W., Rosahl, S., Scheel, D., Llorente, F., Molina, A., Parker, J., Somerville, S., and Schulze-Lefert, P. (2005). Pre- and postinvasion defenses both contribute to nonhost resistance in *Arabidopsis*. Science 310, 1180–1183.

47. Chen, X.L., Yang, J., and Peng, Y.L. (2011). Large-scale insertional mutagenesis in *Magnaporthe oryzae* by *Agrobacterium tumefaciens*-mediated transformation. Methods Mol. Biol. 722, 213–224.

48. Liu, Y. G., Mitsukawa, N., Oosumi, T., and Whittier, R.F. (1995). Efficient isolation and mapping of *Arabidopsis thaliana* T-DNA insert junctions by thermal asymmetric interlaced PCR. Plant J. 8, 457–463.

49. Peñalva, M.A., Lucena-Agell, D., and Arst, H.N. Jr. (2014). Liaison alcaline: Pals entice non-endosomal ESCRTs to the plasma membrane for pH signaling. Curr. Opin. Microbiol. 22, 49–59.

50. Strasser, K., McDonnell, E., Nyaga, C., Wu, M., Wu, S., Almeida, H., Meurs, M.J., Kosseim, L., Powlowski, J., Butler, G. and Tsang, A. (2015). mycoCLAP, the database for characterized lignocellulose-active proteins of fungal origin: resource and text mining curation support. Database (Oxford) 8, 2015.

51. Liu W, Xie S, Zhao X, Chen X, Zheng W, Lu G. Wang Z. (2010) A homeobox gene is essential for conidiogenesis of the rice blast fungus *Magnaporthe oryzae*. Mol Plant Microbe Interact 23(4): 366–375.

52. Tsuji G, Kenmochi Y, Takano Y, Sweigard J, Farrall L, et al. (2000) Novel fungal transcriptional activators, Cmr1p of *Colletotrichum lagenarium* and pig1p of *Magnaporthe grisea*, contain Cys2His2 zinc finger and Zn(II)2Cys6 binuclear cluster DNA-binding motifs and regulate transcription of melanin biosynthesis genes in a developmentally specific manner. Mol Microbiol 38, 940–954.

53. Landraud, P., Chuzeville, S., Billon-Grande, G., Poussereau, N., and Bruel, C. (2013). Adaptation to pH and role of PacC in the rice blast fungus *Magnaporthe oryzae*. PLoS ONE 8, e69236.

54. Alkan, N., Meng, X., Friedlander, G., Reuveni, E., Sukno, S., Sherman, A., Thon, M., Fluhr, R., and Prusky, D. (2013). Global aspects of pacC regulation of pathogenicity genes in *Colletotrichum gloeosporioides* as revealed by transcriptome analysis. Mol. Plant Microbe Interact. 26(11), 1345–1358.

55. Caracuel, Z., Roncero, M.I., Espeso, E.A., Gonzalez-Verdejo, C.I., Garcia-Maceira, F.I., and Di Pietro, A. (2003). The pH signalling transcription factor PacC controls virulence in the plant pathogen Fusarium oxysporum. Mol Microbiol 48, 765–779.

56. Chen, Y., Li, B.Q., Xu, X.D., Zhang, Z.Q., and Tian, S.P. (2018). The pH-responsive PacC transcription factor plays pivotal roles in virulence and patulin biosynthesis in *Penicillium expansum*. Environ. Microbiol. 20, 4063–4078.

57. Rollins, J.A., and Dickman, M.B. (2001). PH signaling in Sclerotinia sclerotiorum: Identification of a pacC/RIM1 Homolog. Appl Environ Microb 67, 75–81.

58. You, B.J., Choquer, M., and Chung, K.R. (2007). The Colletotrichum acutatum gene encoding a putative pH-responsive transcription regulator is a key virulence determinant during fungal pathogenesis on citrus. Mol Plant Microbe Interact. 20, 1149–1160.

59. Lucena-Agell, D, Hervas-Aguilar, A., Munera-Huertas, T., Pougovkina, O., Rudnicka, J., Galindo, A., Tilburn, J., Arst, H.N. Jr., and Peñalva, M.A. (2016). Mutational analysis of the *Aspergillus* ambient pH receptor PalH underscores its potential as a target for antifungal compounds, Mol Microbiol, 101, 982–1002.

60. Li W. and Mitchell A. P. (1997). Proteolytic Activation of Rimlp, a Positive Regulator of Yeast Sporulation and Invasive Growth. Genetics 145, 63–73

61. Futai E., Maeda T., Sorimachi H., Kitamoto K., Ishiura S., Suzuki K. (1999). The protease activity of a calpain-like cysteine protease in Saccharomyces cerevisiae is required for alkaline adaptation and sporulation. Mol Gen Genet 260, 559–568.

62. Espeso, E.A., and Arst, H.N. Jr., (2000). On the mechanism by which alkaline pH prevents expression of an acid-expressed gene. Mol Cell Biol. 20, 3355–3363.

63. Yang, J., Zhao, X.Y., Sun, J., Kang, Z.S., Ding, S.L., Xu, J.R., and Peng, Y.L. (2010) A novel protein Com1 is required for normal conidium morphology and full virulence in *Magnaporthe oryzae*. Mol. Plant Microbe Interact. 23, 112–123.

64. Talbot, N.J., Kershaw, M.J., Wakley, G.E., De Vries, O.M.H., Wessels, J.G.H., and Hamer, J.E. (1996). *MPG1* encodes a fungal hydrophobin involved in surface interactions during infection-related development of *Magnaporthe grisea*. Plant Cell 8, 985–999.

65. Peng, Y.L., and Shishiyama, J. (1988). Temporal sequence of cytological events in rice leaves infected with *Pyricularia oryzae*. Can. J. Bot. 66, 730–735.

66. Xu, J.R., and Hamer, J.E. (1996). MAP kinase and cAMP signaling regulate infection structure formation and pathogenic growth in the rice blast fungus *Magnaporthe grisea*. Genes Dev. 10(21), 2696–2706.

67. Sambrook, J., and Russell, D.W. (2001). Molecular cloning: A laboratory manual. (Cold Spring Harbor, NY: Cold Spring Harbor Laboratory Press).

68. Park, G., Xue, C., Zhao, X., Kim, Y., Orbach, M., Xu, J.R. (2006) Multiple upstream signals converge on the adaptor protein Mst50 in *Magnaporthe grisea*. Plant Cell. 18, 2822–2835.

69. Xue, M., Yang, J., Li, Z., Hu, S., Yao, N., Dean, R.A., Zhao, W., Shen, M., Zhang, H., Li, C., Liu, L., Cao, L., Xu, X., Xing, Y., Hsiang, T., Zhang, Z., Xu, J.R., and Peng, Y.L. (2012). Comparative analysis of the genomes of two field isolates of the rice blast fungus *Magnaporthe oryzae*. PLoS Genet. 8, e1002869.

70. Kim, D., Pertea, G., Trapnell, C., Pimentel, H., Kelley, R., and Salzberg. S.L. (2013). TopHat2: accurate alignment of transcriptomes in the presence of insertions, deletions and gene fusions. Genome Biol. 14(4), R36.

71. Trapnell, C., Williams, B.A., Pertea, G., Mortazavi, A., Kwan, G., van Baren, M.J., Salzberg, S.L., Wold, B.J., and Pachter, L. (2010). Transcript assembly and quantification by RNA-Seq reveals unannotated transcripts and isoform switching during cell differentiation. Nat. Biotechnol. 28, 511–515.

72. Vanheeswijck, R. and Hynes, M.J. (1991). The amdR product and a CCAAT-binding factor bind to adjacent, possibly overlapping DNA sequences in the promoter region of the *Aspergillus nidulans* amdS gene. Nucleic Acids Res. 19, 2655–2660.

